# Antibiotic effects on microbial communities are modulated by resource competition

**DOI:** 10.1101/2022.09.01.506215

**Authors:** Daniel Philip Newton, Po-Yi Ho, Kerwyn Casey Huang

## Abstract

Antibiotic treatment significantly impacts the human gut microbiota, but quantitative understanding of how antibiotics affect community diversity is lacking. Here, we build on classical ecological models of resource competition to investigate community responses to antibiotic-induced species-specific death rates. Our analyses highlight the complex dependence of species coexistence that can arise from the interplay of resource competition and antibiotic activity, independent of other biological mechanisms. We show that resource competition can cause richness to change non-monotonically as antibiotic concentrations are increased. We identified resource competition structures that cause richness to depend on the order of sequential application of antibiotics (non-transitivity), and the emergence of synergistic and antagonistic effects under simultaneous application of multiple antibiotics (non-additivity). These complex behaviors can be prevalent, especially when generalist consumers are targeted. Communities can be prone to either synergism or antagonism, but typically not both, and antagonism is more common. Furthermore, we identify a striking overlap in competition structures that lead to non-transitivity during antibiotic sequences and those that lead to non-additivity during antibiotic combination, suggesting that our analysis is broadly applicable across a wide range of clinically relevant antibiotic treatment schemes. In sum, our results will facilitate the engineering of community dynamics via deleterious agents.

## INTRODUCTION

Antibiotics are cornerstones of modern medicine due to their ability to inhibit the growth of pathogens during infections. Antibiotics are broadly characterized by their inhibitory mechanism as bacteriostatic (growth-halting) or bactericidal (death-causing), and by their spectrum of activity as broad or narrow (Kohanski et al., 2010). The effects of most antibiotics have been predominantly investigated in species monocultures (Andrews, 2001), despite the fact that treatment in a clinical setting inevitably has unintended consequences on the multispecies communities that colonize the human gut (Cani, 2018). Antibiotics can exert collateral damage on gut bacteria (Maier et al., 2021) and reduce gut microbiota diversity (Aranda-Diaz et al., 2022; Maier et al., 2018; Ng et al., 2019), the latter of which has been linked to increased propensity for *Clostridiodes difficile* infection (Hromada et al., 2021; Owens Jr et al., 2008; Schubert et al., 2015) and increased mortality after cancer treatment (Taur et al., 2014). A deeper understanding of the interplay between antibiotic activity and community context could provide mechanisms to ameliorate treatment side effects and to mitigate the growing threat of antibiotic resistance (De Leenheer and Cogan, 2009; Hallinen et al., 2020).

Community dynamics during antibiotic perturbations can be challenging to predict since context can alter antibiotic effects through interspecies interactions, pH modulation, and metabolic transformation (Aranda-Diaz *et al*., 2022; Bottery et al., 2021; de Vos et al., 2017; Zimmermann et al., 2021). The use of a variety of treatment regimens such as variable dosage (De Leenheer and Cogan, 2009), sequential scheduling of multiple compounds (Batra et al., 2021), and concurrent treatment with cocktails (Wood and Cluzel, 2012) further complicates predictions. Moreover, concurrent application of multiple drugs can lead to synergistic or antagonistic effects (Brochado et al., 2018; Xu et al., 2018; Yeh et al., 2009), whereby the degree of killing is greater or less, respectively, than the sum of the drugs individually. Importantly, intrinsic competition for nutrients within a community is a major driver of community dynamics in the absence (Gowda et al., 2022; Ho et al., 2022a; Ho et al., 2022b; Segura Munoz et al., 2022) and presence of antibiotics (Adamowicz et al., 2018; Amor and Gore, 2022). This plethora of phenomena motivates the development of a theoretical approach to untangle the network of interspecies interactions from activity of the antibiotic itself.

To interrogate the potential for emergent effects of antibiotics on community dynamics, we utilized consumer-resource (CR) models in which species growth is governed by nutrient availability (Chesson, 1990). Within CR models, species can coexist only when they occupy distinct resource niches (Marsland et al., 2019; Posfai et al., 2017; Taillefumier et al., 2017); how antibiotic perturbations affect species coexistence remains unclear. Here, we incorporate species-specific death rates into CR models to map the complex behaviors that can arise from the interplay between antibiotic activity and resource competition. We derived a general framework describing how species-specific death rates shift the resource competition landscape, and hence the criteria for coexistence and the consequences for community diversity. Using this framework, we delineated the effects of antibiotic dosage, scheduling, and combination. We found that increasing the degree of targeting of a single species, akin to varying the concentration of a narrow-spectrum antibiotic, can result in non-monotonic changes in richness. In addition, the order of sequential application of multiple antibiotics can qualitatively affect the final community architecture, and treatment with antibiotic combinations can result in synergism or antagonism at the level of community diversity. These phenomena arose solely from resource competition, independent of other biological mechanisms. Importantly, these phenomena were prevalent, suggesting that they are likely to occur in typical communities of gut commensals. Thus, our results suggest that communities can be designed to exploit resource competition for improving therapeutics.

## RESULTS

### Conditions for coexistence in a CR model with antibiotic activity

We explore the effects of antibiotics on community dynamics using an established formulation of a well-mixed CR model of *m* species competing for *p* supplied resources in a chemostat (Posfai *et al*., 2017) described by the equations

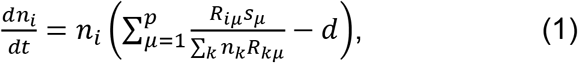

where *n_i_* is the abundance of species *i*. *R_iμ_* is the rate at which species *i* consumes resource *μ, s_μ_* is the supply rate of resource *μ*, and *d* is the dilution rate of the chemostat, which affects each species uniformly (Fig. 1A, Table 1). To model the effects of bacteriostatic antibiotics, we assumed that the consumption rates *R_iμ_* of species *i* decrease by a factor *b_i_*, modifying Eq. 1 as follows:

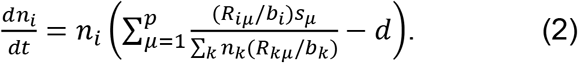

**Figure 1:**
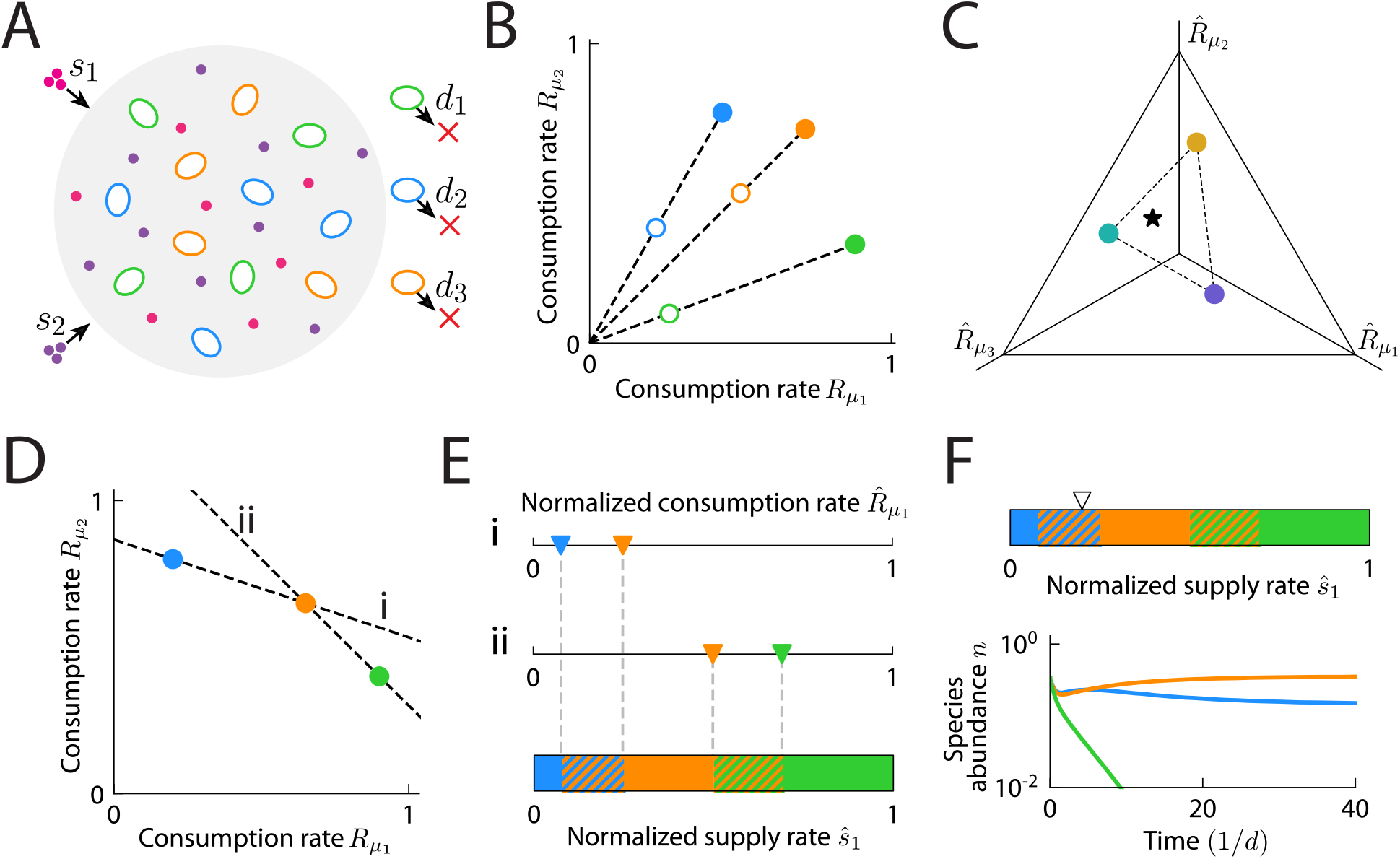
Implementation of antibiotic activity in a CR model. A) Schematic of a well-mixed CR model (Posfai *et al*., 2017) with species-specific death rates *d_i_*. Depicted are cells (colored ovals) from three species competing for two resources (small circles) supplied at rates *s_μ_*. B) The effects of species-specific death rates on coexistence can be determined by reducing the consumption rates *R_iμ_* by *b_i_* = (*d* + *d_i_*)/*d*. The transformed consumption rates (open circles) have reduced enzyme budgets compared to the original consumption rates (filled circles). C) When a community of three species competes for three resources, the consumption rates can be visualized on a simplex representing the hyperplane containing the consumption niches of the three species in the space of rescaled consumption rates. In the case shown, the convex hull of the species consumption rate vectors (dashed line) encloses the point representing the normalized resource supply rates (star), hence all species coexist (Posfai *et al*., 2017). D) Example in which two possible hyperplanes (dashed lines, i and ii) dictate the conditions for coexistence of three species (colored circles) competing for two resources. E) Along each hyperplane, a pair of species coexists if the normalized resource supply rates *ŝ_μ_* lie between the rescaled consumption rates (colored triangles) of the species pair (hatched multicolored regions). Otherwise, only the species with 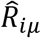 closest to *ŝ_μ_* persists (solid regions). F) Species dynamics (bottom) for the community shown in (D) for one case of supply rates (triangle, top).

**Table 1:** Definitions of key variables and terms.

| Symbol | Definition |
| --- | --- |
| $m$ | Number of species initially present |
| $p$ | Number of types of supplied resources |
| $n_i(t)$ | Abundance of species $i$ at time $t$ |
| $n_i^*$ | Steady state abundance of species $i$ |
| $\rho$ | Richness, or number of species with nonzero abundance at steady state |
| $R_{i\mu}$ | Rate at which species $i$ consumes resource $\mu$ |
| $s_\mu$ | Supply rate of resource $\mu$ |
| $d$ | Dilution rate of the chemostat |
| $b_i$ | Species-specific reduction factor of $R_{i\mu}$ values corresponding to species $i$, interpreted as the susceptibility of species $i$ to antibiotics |
| $d_i$ | Death rate of species $i$ |
| $E_i$ | Enzyme budget of species $i$, where $E_i = \sum_{\mu}^p s_{\mu}$ |
| $S$ | Total resource supply rate (in general, assumed to be $S = 1$) |
| $c_{\mu}^*$ | Steady-state concentration of resource $\mu$ |
| $\hat{s}_{\mu}$ | Normalized supply rate of resource $\mu$, where $\hat{s}_{\mu} = (d/S)s_{\mu}$ |
| $\hat{R}_{i\mu}$ | Rescaled rate at which species $i$ consumes resource $\mu$, where $\hat{R}_{i\mu} = c_{\mu}^* R_{i\mu}$ |
| coexistence region size or size of convex hull | Fraction of supply rates that lead to coexistence |
| specialist/generalist consumer | Species that consumes one/all resources at nonzero rates |
| “more” specialist/generalist consumer | Species that has a more/less even distribution of consumption rates |
| $c$ | Factor that scales all values of $d_i$ equally to mimic the concentration of an antibiotic |
| $D_{Ti}$ | Magnitude of the difference between $\hat{R}_i$ and $\hat{s}$, where species $i$ is targeted by antibiotics |
| $D_T$ | Magnitude of the difference between $\hat{R}_i$ and $\hat{R}_j$, where $i$ and $j$ are the two species targeted by antibiotics |
| $D_N$ | Magnitude of the difference between $\hat{R}_i$ and $\hat{s}$, where $i$ is the non-targeted species |
| $d_{\min}$ | Minimum value of $d_i$ among all nonzero species-specific death rates $d_i$ during antibiotic perturbation |
| $r_i^*$ | Relative abundance of species $i$ at steady state |
| $N_{\text{eff}}$ | Effective number of species at steady state ($N_{\text{eff}} = \exp(-\sum_i r_i^* \ln(r_i^*))$), where $r_i^*$ is the relative abundance of species $i$ at steady state |

To model the effects of bactericidal antibiotics, we assumed that species *i* experiences death at rate *d_i_* in addition to the effects of dilution (Fig. 1A), modifying Eq. 1 as follows:

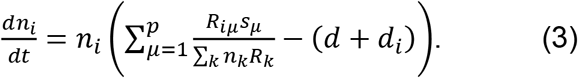

Importantly, when *b_i_* = (*d* + *d_i_*)/*d*, Eq. 2 and 3 are identical up to rescalings of time, species abundances, and resource consumption rates (Supplemental Text). Therefore, within this model, the effects of antibiotic activity on species coexistence can be understood as a reduction of the enzyme budget of species *i, E_i_* = ∑*_μ_R_iμ_*, by a species-specific factor *b_i_* regardless of antibiotic mechanism (Fig. 1B). Steady-state species abundances are also typically similar between bactericidal and bacteriostatic antibiotic activity given the transformation described above, particularly for communities with larger overlap in consumption niche (Fig. S1A,B), suggesting that antibiotic mechanism of action may be less impactful on community composition than the nutrient niches of the targeted species.

It was previously shown that within the chemostat CR model described by Eq. 1, a set of species *i* with the same enzyme budgets (*E_i_* = *E*) will coexist if the convex hull of their consumption rates contains the normalized resource supply rates (*E/S*)*s_μ_* (Fig. 1C), where *S* = ∑*_μ_s_μ_* is the total resource supply rate (Posfai *et al*., 2017). Motivated by the implementation of antibiotic activity described above, we found that we could generalize this coexistence rule to species-specific enzyme budgets *E_i_* and *d_i_* = 0 without loss of generality as follows. For a set of *m* species, a subset of *m′* species will coexist if their consumption rates lie on a hyperplane defined by 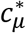 such that 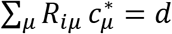 for all species *i* in the subset, and if the hyperplane satisfies the following requirements: 1) all axis-intercepts 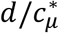 are positive and finite (so that the steady state resource concentrations 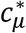 are positive and finite); 2) no species have consumption rates that lie on the side of the hyperplane away from the origin, i.e., 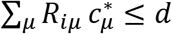 for all species *i* (since a species whose consumption rates lie above the hyperplane has a large enough enzyme budget that it will drive species on the hyperplane extinct); and 3) the normalized resource supply rates *ŝ_μ_* = *s_μ_*(*d/S*) lie within the convex hull of the rescaled consumption rates 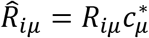 of the *m′* species (namely, the resource supply rates must lie within a region of shared consumption niche among all coexisting species) (Supplemental Text). In an example community with three species competing for two resources, there are two hyperplanes (lines) that satisfy the first two conditions (Fig. 1D), and along each line, the species that will persist are determined according to the third condition (Fig. 1 E,F). Unless otherwise specified, we will work in the space of rescaled consumption rates and normalized supply rates with no loss of generality (Supplemental Text), and we assume *d* = *S* = 1. As presented in the sections below, these rules produce some intuitive behaviors as well as complex behaviors emerging from the interplay between species-specific antibiotic activity and resource competition, each of which can inform the interpretation of experiments.

### Antibiotic treatment can promote coexistence in resource competition regimes involving generalists

To interrogate how antibiotic activity affects species coexistence, we first applied the coexistence conditions in Fig. 1 to analyze the minimal scenario of two species competing for two resources, *m* = *p* = 2. We considered four qualitatively distinct types of communities: two specialist consumers (each with only one nonzero consumption rate) with no niche overlap, one generalist (with two nonzero consumption rates) and one specialist, a pair of generalists with preference for distinct resources, and a pair of generalists with preference for the same resource (Fig. 2A). Naively, one might expect that increasing the death rate of a species would generally decrease the probability that it can persist in a community, and hence decrease the proportion of supply rates that lead to coexistence (defined by the size of the convex hull or coexistence region). For two generalists with preference for distinct resources, as the ratio of their enzyme budgets changed due to antibiotic perturbation, the coexistence region size indeed decreased (Fig. 2A, red). However, for the trivial case of two specialists, there is no competition and hence the two species coexisted regardless of death rates (Fig. 2A, purple). For a generalist and a specialist, decreasing the enzyme budget of the generalist increased the coexistence region size (Fig. 2A, green), indicating that antibiotic activity targeting a generalist can promote coexistence. For two generalists with preference for the same resource, coexistence region size can exhibit non-monotonic dependence on enzyme budgets (Fig. 2A, yellow).

**Figure 2:**
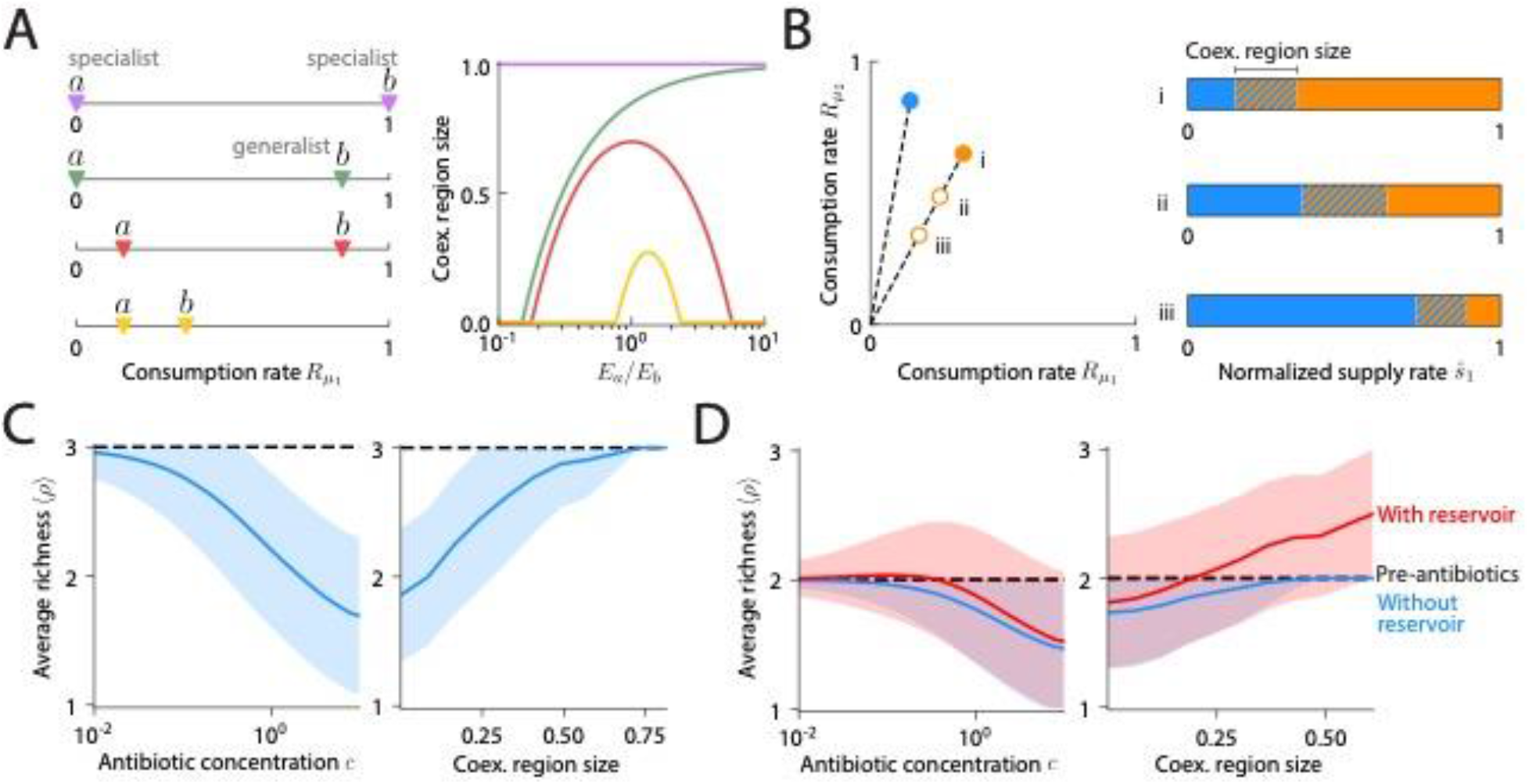
Resource competition results in complex dependence of species coexistence on antibiotic activity. A) Effect of antibiotic activity on two species (*a* and *b*) competing for two resources. (Left) Four types of communities with qualitatively different consumption niches were considered, including a specialist and a generalist (green), two specialists (purple), two symmetrically distributed generalists (red), and two asymmetrically distributed generalists (yellow). (Right) The fraction of supply rates that lead to coexistence (coexistence region size) can depend non-monotonically on the relative enzyme budget (*E_a_/E_b_* of species *a* and *b* on the left). B) Example community consisting of a non-targeted species (blue) and a targeted species (orange) that was initially more of a generalist (more equal consumption of both resources). (Left) Antibiotic activity resulted in decreased enzyme budget (empty circles). (Right) Map of coexistence regions as in Fig. 1E for the scenarios on the left. C) Average richness decreases with increasing antibiotic concentration *c* (left) and decreasing coexistence region size (right). Simulations of randomly drawn communities of three species competing for three resources (*m* = *p* = 3) with consumption rates 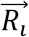 sampled uniformly from the unit simplex and all resources supplied at equal rates. Average richness calculated across all communities with richness of 3 before antibiotic perturbation. Antibiotic concentration *c* scales the death rates, i.e., *d_i_* → *cd_i_*. D) A reservoir of species can lead to an increase in average richness during antibiotic perturbation. As in (C), but for communities with richness of 2 before antibiotic perturbation without (blue) or with a re-seeding pool of species to transiently repopulate the initially distinct species during antibiotic perturbation (red).

These non-monotonic behaviors arise because antibiotic activity affects the shape of the coexistence region rather than simply rescaling its size (Fig. 2B). In the case of two generalists, when the more generalist of the two species (the one with more equal consumption of both resources) was targeted (Fig. 2B, i), its consumption niche was encroached upon by the non-targeted species (Fig. 2B, ii), effectively making the non-targeted species more of a generalist. As a result, the bounds of the coexistence region shifted toward the consumption niche of the targeted species since supply rates must be closer to the consumption rates of the targeted species to enable its coexistence (Fig. S2). Thus, the coexistence region size initially increased with death rate and reached its maximum when the remapped boundaries were symmetric about the point of equal supply rates (Fig. 2B, ii). When the death rate of the targeted species was increased further (Fig. 2B, iii), expansion of the consumption niche of the non-targeted species and the concomitant shrinkage of the consumption niche of the targeted species led to an overall decrease in coexistence region size (Fig. 2B, iii). Steady-state community evenness in species abundance (measured as the exponential of the Shannon diversity index (Jost, 2006)) was similarly non-monotonic for this scenario (Fig. S1C), suggesting that antibiotic perturbations have similar effects on richness *ρ* and evenness. This analysis highlights the potential for complex behavior involving species coexistence when a generalist is targeted.

Despite the example above, we hypothesized that antibiotic activity should generally decrease richness when averaged across communities. To test this hypothesis, we investigated communities with three species competing for three resources (*m* = *p* = 3) with consumption rates 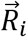 sampled uniformly from the unit simplex of equal enzyme budgets (i.e., 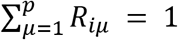 with equal probability of sampling all vectors that satisfy this constraint). Species-specific death rates were drawn uniformly from the unit simplex for each community, and all resources were supplied at equal rates. For communities in which all three species coexisted prior to antibiotic perturbation, the average richness across all communities indeed decreased following the implementation of species-specific death rates, and the decrease in richness was larger for communities with smaller coexistence regions (Fig. 2C, right). Consistent with this trend, mimicking an increase in antibiotic concentrations for each community by multiplying all species-specific death rates by a factor *c* caused average richness to decrease monotonically (Fig. 2C, left).

For the decrease in average richness described above, all starting communities had maximal richness. To consider a scenario in which richness has more potential to rise upon antibiotic treatment, we focused on communities in which only two of the three species coexisted prior to antibiotic perturbation, and allowed for an external reservoir of species to transiently repopulate the initially extinct species during the antibiotic perturbation, as has been observed experimentally (Ng *et al*., 2019). Now, the average richness reached a maximum >2 for communities with small antibiotic concentrations or intermediate coexistence regions (Fig. 2D). Thus, non-monotonic richness behavior upon antibiotic treatment can be a prevalent feature of communities in the presence of a re-seeding pool of species.

### Increasing death rate can lead to highly non-monotonic changes in richness

Our simulations of randomly drawn communities showed that richness can increase upon antibiotic treatment in some cases (Fig. 2C). To investigate this behavior in more detail, we considered a community with *m* = *p* = 3 in which only one species (blue) persisted at steady state prior to antibiotic perturbation (Fig. 3A,B). As the death rate of this species was increased, in the presence of a re-seeding reservoir of all three species, the other two species (orange and green) were able to coexist with the targeted species (blue) during low levels of antibiotic perturbation (Fig. 3B, *d*_1_ = 0.5), analogous to the initial increase in coexistence region size when the more generalist species was targeted in a two-member community (Fig. 2B). As the death rate of the targeted species was increased further, the rescaled consumption rates indicated that the targeted species became more specialized (i.e., moved away from the supplied resource point at the center of the simplex), whereas the other two species became more like generalists due to the relative increase in their enzyme budgets compared to that of the targeted species. The perturbed convex hull shifted toward the niche of the targeted species (Fig. 3A, S2) and in doing so, transited through six coexistence states (Fig. 3B), including two where all three species coexisted (Fig. 3B, *d*_1_ = 0.5, *d*_1_ = 8), until the targeted species eventually became extinct at large enough death rates (Fig. 3B, *d*_1_ = 10). This example demonstrates that increasing antibiotic concentration can lead to numerous, non-monotonic richness changes representing different coexistence states.

**Figure 3:**
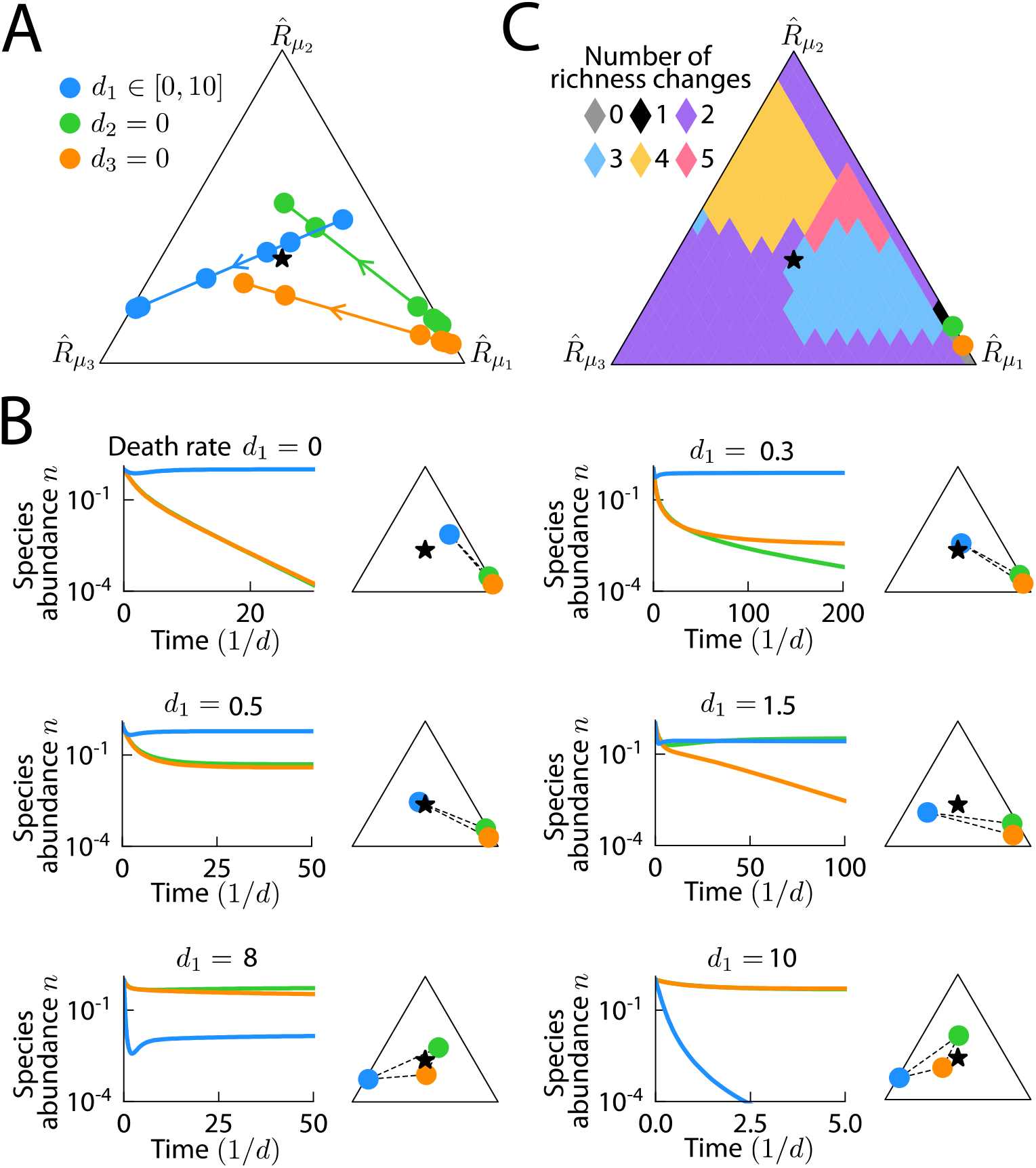
Richness can change non-monotonically with increasing death rate. A) Trajectories of rescaled consumption rates as the death rate of the targeted species (blue) is increased. Arrow denotes the direction of increasing death rate. Circles mark the six death rates shown in (B). With increasing death rate, the targeted species became more specialized, while the non-targeted species became more of a generalist and closer to the supplied resource point at the center of the simplex. B) Population dynamics at the six death rates marked in (A). (*d*_1_ = 0) In the absence of antibiotic targeting, only the blue species persisted. (*d*_1_ = 0.3) For low death rates of the blue species, the orange species was able to coexist. (*d*_1_ = 0.5) There was a range of death rates that allowed for coexistence of all three species. (*d*_1_ = 1.5) As the death rate was increased further, the orange species went extinct. (*d*_1_ = 8) As the death rate increased even further, coexistence of all three species was again realized. (*d*_1_ = 10) Finally, at sufficiently high death rate the blue species went extinct. C) Number of richness changes until the extinction of the targeted species as its consumption niche was varied across the simplex, while the consumption niches of the non-targeted species were fixed as in (A).

To quantify the prevalence of non-monotonic richness changes in response to increasing antibiotic concentration, we fixed the resource consumption rates of the two non-targeted species as in Fig. 3A and varied the consumption rates of the targeted species throughout the simplex. For each set of consumption rates, we calculated the number of changes in community richness as the death rate of the targeted species was increased from zero until the targeted species went extinct. As long as the targeted species was able to coexist prior to antibiotic perturbation, which was true for almost all of the simplex in this example, there were at least two richness changes (Fig. 3C), corresponding to a non-targeted species emerging from extinction into coexistence followed by the targeted species going extinct. Moreover, when the consumption niche of the targeted species was biased against the resource that was least preferred by the non-targeted species (resource 3 in Fig. 3C), indicating more competition, the number of richness changes was typically larger, with a maximum value of five occurring when the targeted species consumed its two preferred resources at approximately equal rates (Fig. 3C). Taken together, these results demonstrate that non-monotonic behavior is likely to be prevalent across resource competition landscapes.

### Non-transitive effects during sequential antibiotic treatments typically arise from promotion of antibiotic-induced extinctions by resource competition

In cases involving targeting of a pathogen, antibiotics are commonly administered sequentially to reduce the emergence of antibiotic resistance in the pathogen (Batra *et al*., 2021). This sequential treatment can have nonintuitive effects on commensal members of the microbiota. To predict the effects of sequential treatment in a community context, we asked whether the final richness in our model is dependent on the sequence in which two antibiotics are sequentially applied (transitivity). We first examined a community of three coexisting species and simulated the sequential application of two narrow-spectrum antibiotics that each target one of the three species (Fig. 4A). For each antibiotic, we simulated the population dynamics until steady state was reached. In this example, the blue species was driven extinct when it was targeted first (Fig. 4B, top left). Next, the antibiotic targeting the blue species was removed. The orange species was then targeted and became extinct as the green species encroached on its niche (Fig. 4B, top right). Thus, this treatment sequence eventually led to the presence of only one species (Fig. 4C, top).

**Figure 4:**
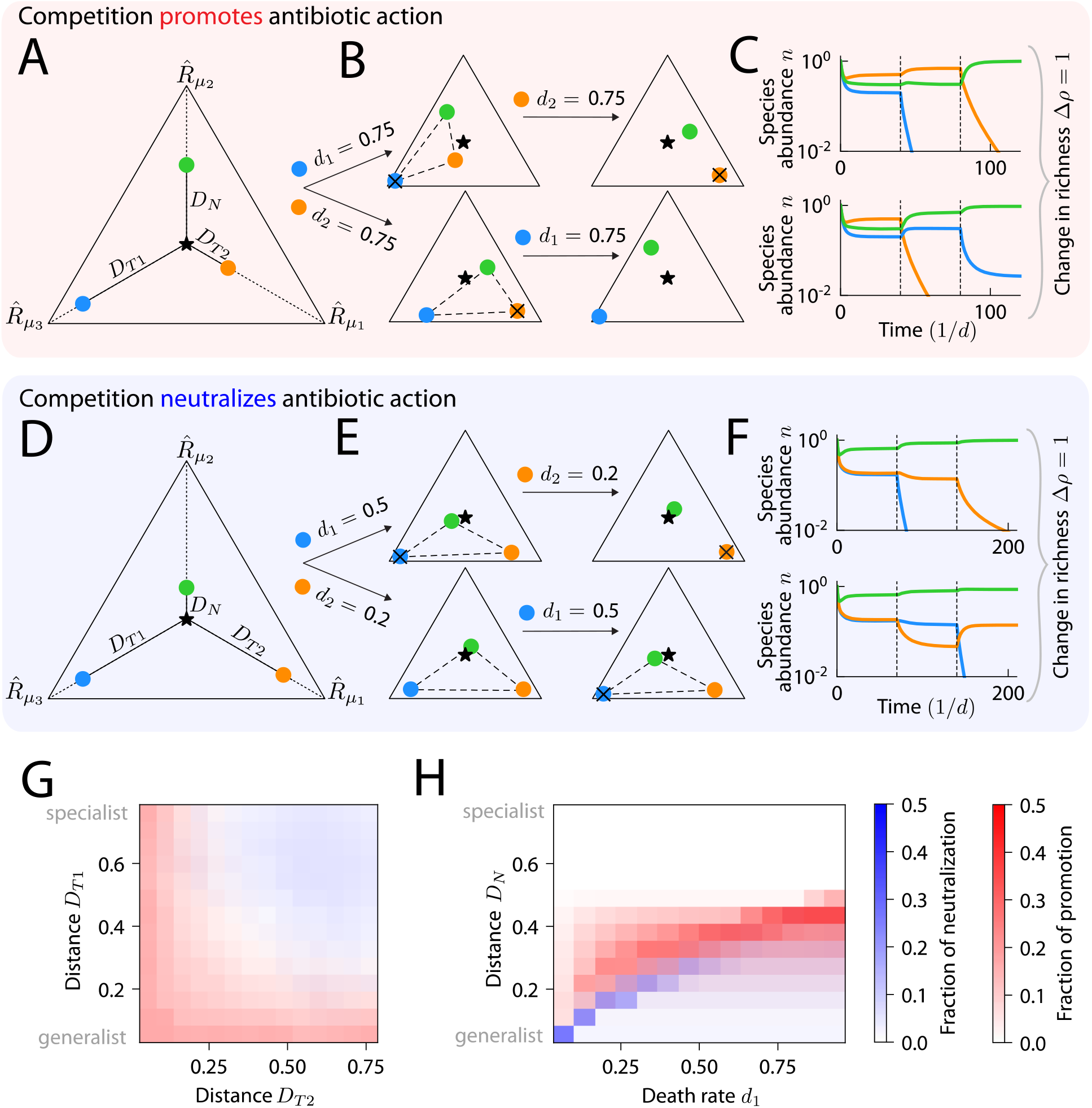
Final coexistence is dependent on the sequence of antibiotic treatment. A) An example community of three species (colored circles) for which the sequence of narrow-spectrum antibiotics can affect the final richness. The orange species promoted the action of the antibiotic that targets the blue species, causing the final richness to be non-transitive to the sequence of antibiotic treatment. B) (Top left) Remapping of the consumption niche of all species when the death rate of the blue species was increased. The blue species went extinct, as the remapped convex hull does not enclose the supplied resource point. (Top right) Increasing the death rate of the orange species after the extinction of the blue species led to its extinction. (Bottom row) Same as top row, but the sequence of sequential antibiotics was reversed. Now, the blue and green species coexisted after the two treatments, thus Δ*ρ* = 1. C) Population dynamics were simulated until steady state for the scenarios in (B): (top) first with zero death rates, then with nonzero death rate for the blue species, then with nonzero death rate for the orange species (and zero death rate for the now extinct blue species); (bottom) with the reversed sequence of death rates. D) An example community with nonzero Δ*ρ* due to the blue species neutralizing the action of the antibiotic that targets the orange species. E) Like (B) but for the community in (D). F) Like (C) but for the community in (D). G) Prevalence of non-transitivity via promotion (red) or neutralization (blue) (or some combination of the two shown by overlapping red and blue) as a function of the resource competition structure. Δ*ρ* was calculated and the mechanism of non-transitivity was determined for communities of the form in (A) and (D) across parameters (*D*_*T*1_, *D*_*T*2_, *D_N_, d*_1*t*_ *d*_2_), for death rates *d*_1_ ∈ (0,1), *d*_2_ ∈ (0,1) and *D*_*T*1_, *D*_*T*2_, and *D_N_* across their entire domains. The fraction of promotion and neutralization were averaged across all combinations *D_N_, d*_1_, and *d*_2_. Non-transitivity due to promotion (neutralization) was more likely for low (high) *D*_*T*1_ and low (high) *D*_*T*2_. H) Same as (G) but averaged across all *D*_*T*1_, *D*_*T*2_, and *d*_2_ (or equivalently, *d*_1_ if *d*_2_ was shown on the x-axis). The fraction of neutralization was large when *D_N_* and *d*_1_ were small, and the fraction of promotion was largest when *D_N_* and *d*_1_ were increased.

However, when the antibiotic treatment sequence was reversed, the community reached a distinct state. When the orange species was targeted first, it went extinct (Fig. 4B, bottom left). Next, when the blue species was targeted, it was not outcompeted by the green species (Fig. 4B, bottom right) and the two coexisted (Fig. 4C, bottom). Thus, the reverse sequence of treatment qualitatively altered the final coexistence (Fig. 4C, bottom) and the absolute value of the richness difference between the two sequences Δ*ρ* was 1 (non-transitivity, Fig. 4C). For this community, the extinction of the blue species was dependent on competition between the orange and blue species (Fig. 4B), hence an antibiotic that eliminated the orange species allowed the blue species to coexist with the green species even when the blue species was later targeted (Fig. 4B, bottom). That is, the mechanism leading to non-transitivity was competition promoting the action of the antibiotic that targets the blue species (Fig. 4B, top left, bottom right), while the antibiotic that targets the orange species caused the extinction of the orange species regardless of community context (Fig. 4B, top right, bottom left). Another scenario that can result in non-transitivity is competition neutralizing the action of one antibiotic but not the other. In an example of neutralization (Fig. 4D-F), the antibiotic that targets the orange species only caused extinction of the orange species when the blue species was absent (Fig. 4E, bottom left and Fig. 4E, top right). Because the antibiotic that targets the orange species was neutralized by competition while the antibiotic that targets the blue species caused the blue species to go extinct regardless of community context (Fig. 4E, top left and Fig. 4E, bottom right), Δ*ρ* = 1 (Fig. 4F).

To explore the conditions under which the final richness depends on antibiotic sequencing, we calculated Δ*ρ* for the community in Fig. 4A across a five-dimensional parameter space: the distances 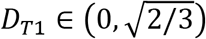 and 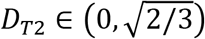 from the supplied resource point to the consumption rates of the two targeted species, the distance between the non-targeted species and the supplied resource point 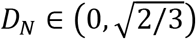 (Fig. 4A), and the death rates of the two targeted species, *d*_1_ ∈ (0, 1), *d*_2_ ∈ (0, 1). We varied each of these five parameters independently across their respective domains, calculating Δ*ρ* for all combinations of parameter values. In all cases, Δ*ρ* was 0 or 1, with Δ*ρ* = 1 when the final richness after an antibiotic sequence was 2 and the final richness after the reverse sequence was 1 (or vice versa); if the final richness after an antibiotic sequence is 3, then the final richness after the reverse sequence must also be 3, and thus Δ*ρ* = 0. This behavior occurs when *D_N_* is sufficiently large (*D_N_* ≳ 0.5), so that the non-targeted species is a specialist for its unique resource and hence cannot outcompete any other species in pairwise competition. As a result, the richness after any antibiotic sequence was ≥ 2 (Fig. S3), hence Δ*ρ* was always 0.

We found that all occurrences of nonzero Δ*ρ* could be classified by resource competition either promoting (Fig. 4A-C) or neutralizing (Fig. 4D-F) antibiotic action for one of the two antibiotics. Neutralization was less common than promotion, with neutralization and promotion accounting for 23.1% and 76.9%, respectively, of nonzero Δ*ρ* cases. Resource competition structures strongly affected the prevalence of non-transitivity by promotion or neutralization (Fig. 4G,H). On average across all values of other parameters, promotion was most likely to occur when *D*_*T*1_ or *D*_*T*2_ was < 0.2 (Fig. 4G, red), in which case one of the targeted species is a generalist and thus can outcompete the other targeted species to extinction (Fig. 4B, top left). Furthermore, promotion was more likely for intermediate *D_N_*~0.4 such that the non-targeted species could outcompete one of the targeted species during pairwise competition but not the other (Fig. 4B). Conversely, neutralization was most likely to occur when both *D*_*T*1_ and *D*_*T*2_ were > 0.4 (Fig. 4G, blue), in which case the targeted species cannot outcompete each other during antibiotics since they are specialists with distinct niches. Instead, extinction occurred because the non-targeted species was a generalist that could outcompete one of the targeted species in pairwise competition (*D_N_* ≲ 0.3, Fig. 4H, blue). For both promotion and neutralization, we found that there must be a balance between *D_N_* and the death rates such that resource competition promotes or neutralizes the action of only one of the antibiotics but not the other (Fig. 4H). In sum, these simulations show that resource competition structures can be prone to non-transitivity under sequential treatment via either promotion or neutralization of antibiotic activity (Fig. 4G,H).

### Antagonistic effects on community richness due to antibiotic combinations are more common than synergism

In addition to dosage and scheduling, another common strategy for treating infections is the simultaneous use a cocktail of multiple antibiotics to avoid the emergence of resistance. An important consideration when choosing antibiotics to act on individual species is the potential for synergistic or antagonistic effects (Baym et al., 2016; Brochado *et al*., 2018; Torella et al., 2010). To investigate the extent of interactions among antibiotics in a community context, we considered three species with equal enzyme budgets competing for three resources (Fig. 5A) with symmetry of the community under swapping of the two targeted species (*D*_*T*1_ = *D*_*T*2_) and the antibiotics that target them (*d*_1_ = *d*_2_). When either one of the targeted species was individually subjected to a sufficiently large death rate, it was driven extinct (Fig. 5B), as expected. However, when the death rates of both targeted species were increased simultaneously, all species were able to coexist (Fig. 5C). The changes in enzyme budgets due to both antibiotics effectively nullified each other, returning the consumption rate vectors to locations near their original locations in the simplex (Fig. 5C), hence neither of the targeted species gained a competitive advantage over the other. Although the non-targeted species (green) became more of a generalist under the activity of both antibiotics, it was initially sufficiently specialized that the remapped convex hull still included the supplied resource point. Thus, for this community, the combination of antibiotics resulted in antagonism.

**Figure 5:**
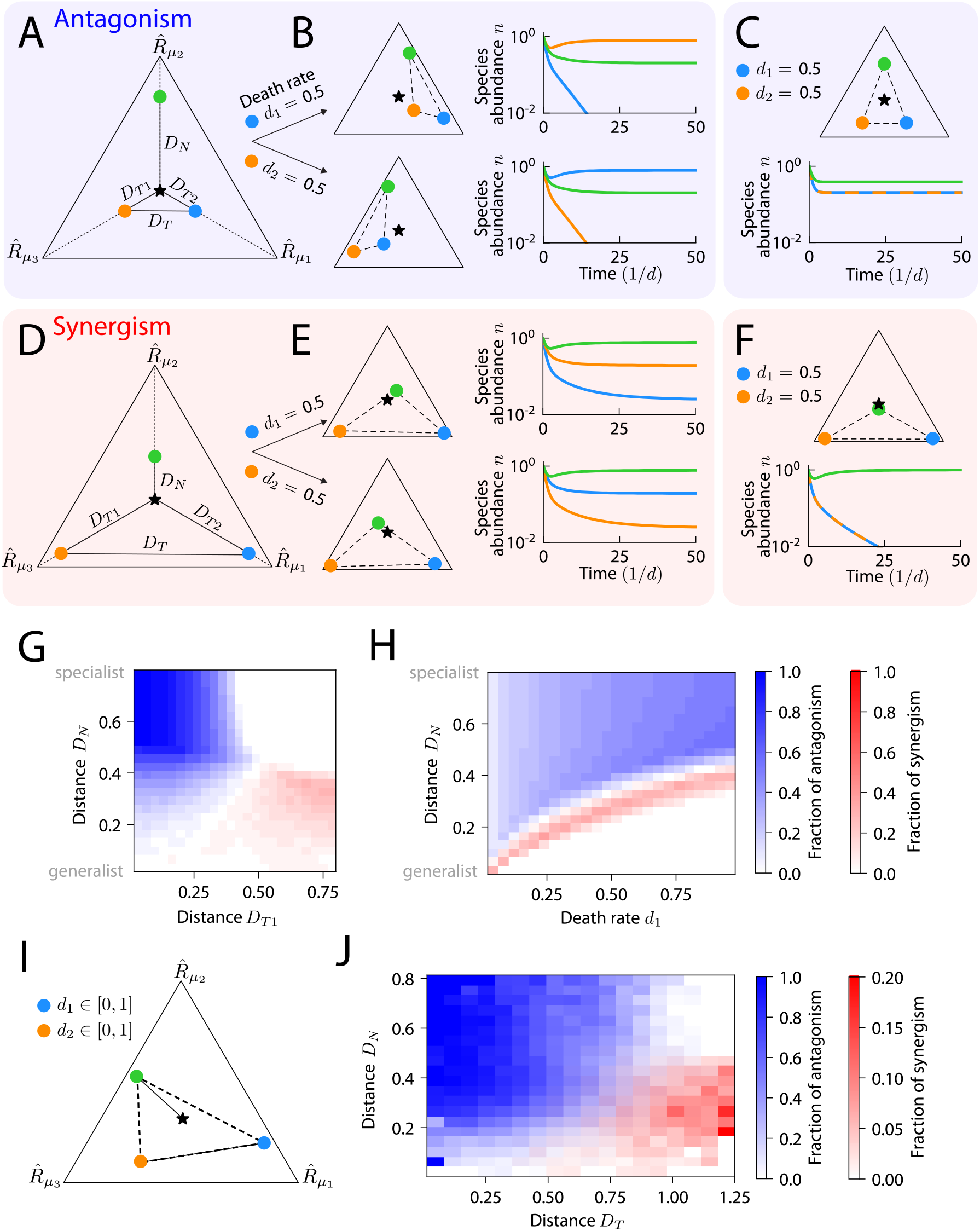
Resource competition can lead to non-additive effects during simultaneous application of two antibiotics, with antagonism more likely than synergism. A) A community for which antibiotic targeting leads to antagonism. Before antibiotic treatment, all three species coexisted, as shown by the simplex of the resource consumption rates of three species (colored circles) along with the supply rates of three resources (star) with distances *D_N_, D*_*T*1_, *D*_*T*2_ as in Fig. 4A, as well as the distance between the two targeted species *D_T_*. B) (Top) When the death rate of the blue species in (A) was increased, remapping of the convex hull resulted in extinction of the blue species. (Bottom) When the death rate of the orange species was increased, the orange species similarly went extinct. C) When both species in (A) were targeted simultaneously, competition between the targeted species was relieved. (Top) Remapping due to both antibiotics led to only small changes in the consumption rate vectors compared with each antibiotic alone, as compared to unperturbed community in (A). (Bottom) Coexistence of all three species was restored, representing antagonism. D) Like (A), but showing a community for which antibiotic targeting led to synergism. E) Targeting of the blue (top) or orange (bottom) species in (D) alone preserved coexistence of all three species. F) When both species in (D) were targeted simultaneously, the non-targeted species (green) outcompeted the targeted species, since the remapped convex hull no longer contained the supplied resource point. The targeted species became extinct, representing synergism. G) The fraction of synergism (red) and antagonism (blue) for communities of the form in (A) and (D) (i.e., *D*_*T*1_ = *D*_*T*2_ and *d*_1_ = *d*_2_ ∈ (0,1)), averaged across all combinations of *D*_*T*2_, *D_N_*, and *d*_2_. Synergism was more likely for high *D*_*T*1_ and low *D_N_*, and vice versa for antagonism. H) Same as (G) but averaged across all values of *D*_*T*1_, *D*_*T*2_, and *d*_2_. The value of *D_N_* at which the fraction of synergism peaked increased as a function of *d*_1_, and antagonism was likely for above this value. I) Schematic of communities with consumption rate vectors drawn uniformly from the simplex, both death rates drawn independently from a uniform distribution between 0 and 1, and resources supplied at equal rates. *D_N_* and *D_T_* are labeled for a random community. J) Of the communities in (I) without symmetry in the consumption rates of the targeted species, a smaller fraction exhibited synergism than antagonism, and the likelihood of either form of non-additivity depended strongly on *D_N_* and *D_T_*, analogous to communities with symmetry as in (G).

Next, we considered a community with larger *D*_*T*1_ = *D*_*T*2_ and with the non-targeted species (green) more of a generalist than in the example above that exhibited antagonism (Fig. 5D). When one of the species was targeted individually, it became more specialized and the non-targeted species moved toward the consumption niche of the targeted species, but the remapped convex hull still enclosed the supplied resource point, resulting in coexistence of all three species (Fig. 5E). However, when both species were simultaneously targeted, the non-targeted species became more competitive against the targeted species and its remapped consumption rates moved past the supplied resource point, allowing it to drive both targeted species extinct (Fig. 5F). The cumulative effect of targeting both species thus resulted in synergism.

To quantify the prevalence of synergism and antagonism across community structures, we varied the distance between the consumption rates of the two targeted species *D_T_* by varying *D*_*T*1_ = *D*_*T*2_, the distance between the non-targeted species and the supplied resource point *D_N_*, and the death rate of both targeted species during antibiotics *d*_1_ = *d*_2_ ∈ (0,1). We varied all three parameters (*D_N_, D*_*T*1_, *d*_1_) independently and calculated the fraction of communities that exhibited synergism or antagonism as a function of two parameters, averaging over the other parameter. Antagonism occurred most often when the niches of the targeted species were somewhat similar (*D_T_* ≲ 0.4) and the non-targeted species was sufficiently specialized (*D_N_* ≳ 0.4), and synergism occurred in the opposite regime (Fig. 5G). This behavior is consistent with the example communities discussed above: for antagonism, each targeted species must be driven extinct when targeted alone (*D_T_* ≲ 0.4) and the non-targeted species must not outcompete the two targeted species when both are simultaneously targeted (*D_N_* ≳ 0.4), and vice versa for synergism. Increasing the death rates of the targeted species commensurately (*d*_1_ = *d*_2_) increased the minimum *D_N_* required for synergism and antagonism (Fig. 5H), as expected.

To investigate whether these trends generalized, we generated communities with consumption rate vectors sampled uniformly across the simplex without enforcing any symmetry (Fig. 5I), sampled the two death rates independently from a uniform distribution between 0 and 1, and supplied all three resources at equal rates. Similar to the trends observed with symmetry (Fig. 5G,H), communities with smaller *D_N_* and larger *D_T_* were more likely to exhibit synergism (Fig. 5J, red) and communities with larger *D_N_* and smaller *D_T_* were more likely to exhibit antagonism (Fig. 5J, blue). Moreover, to exhibit synergism, communities with larger *D_N_* required a larger value of the smaller of the two death rates *d*_min_ (Fig. S4). This condition enables the non-targeted species to outcompete the targeted species, which either requires that the non-targeted species is a generalist that is close to outcompeting the other species before antibiotic perturbation or that the antibiotic perturbation is sufficiently large. Similarly, communities with larger *d*_min_ required a larger *D_N_* to exhibit antagonism (Fig. S4), generalizing the observation that increasing the death rate of the targeted species in the symmetric scenario in Fig. 5A increased the minimum *D_N_* required for antagonism (Fig. 5H). These simulations also demonstrated that communities can be prone to synergistic and antagonistic effects due to resource competition alone during antibiotic combinations, but typically not both (Fig. 5G,H,J), and that antagonism is more common (Fig. 5J).

### Overlap in mechanisms underlying non-transitivity and non-additivity

Since resource competition structures can be prone to neutralizing or promoting antibiotic action during sequential treatment (non-transitivity, Fig. 4) and to synergism or antagonism during simultaneous treatment (non-additivity, Fig. 5), we wondered to what extent the underlying mechanisms were connected. For communities parametrized by *D_N_*, *D*_*T*1_, and *D*_*T*2_, structures prone to neutralization and promotion (Fig. 4H) were approximate subsets of those prone to synergism and antagonism (Fig. 5H), respectively. The same trends generalized to scenarios with randomly drawn structures and death rates (Fig. 5I): Of 100,000 communities, non-transitivity due to promotion and neutralization occurred in 10,482 and 647, respectively, and non-additivity due to antagonism and synergism occurred in 37,782 and 1,156 cases, respectively. Strikingly, 100% of the communities exhibiting non-transitivity due to promotion also exhibited non-additivity due to antagonism, and >97% of the scenarios exhibiting neutralization also exhibited synergism (Table 2). These findings highlight the significant overlap in mechanisms underlying non-additivity and non-transitivity.

**Table 2:** Cross-occurrences of non-transitivity during antibiotic sequences and non-additivity during antibiotic combinations in 100,000 random communities.

| Behavior | Additivity | Antagonism | Synergism | Total |
| --- | --- | --- | --- | --- |
| Transitivity (i.e., $\Delta\rho = 0$) | 61,050 | 27,293 | 528 | 88,871 |
| Nonzero $\Delta\rho$ due to *promotion* | 0 | 10,482 | 0 | 10,482 |
| Nonzero $\Delta\rho$ due to *neutralization* | 12 | 7 | 628 | 647 |
| **Total** | 61,062 | 37,782 | 1,156 | 100,000 |

## DISCUSSION

The ability to predict microbial community dynamics after antibiotic treatment would be a powerful tool for minimizing unintended consequences of treatment. Motivated by growing evidence that resource competition plays a dominant role in shaping microbial community dynamics both without and during antibiotic perturbation, we introduced species-specific death rates to a CR model and analyzed community responses across a wide range of clinically relevant antibiotic treatment strategies. We found that increasing antibiotic intensity can lead to non-monotonic richness changes in the presence of a re-seeding reservoir due to changes in the competitive landscape (Fig. 3), and that the final community can differ qualitatively when the sequence of antibiotic application is reversed (Fig. 4). Furthermore, we quantified properties of resource competition landscapes that give rise to antibiotic synergism or antagonism (Fig. 5), suggesting that the effects of antibiotic perturbation are generally dependent on the metabolic properties of the exposed community. Intriguingly, we find that antagonism emerges in communities with symmetry between the consumption rates of the two targeted species when the non-targeted species is sufficiently specialized (Fig. 5A); this scenario mimics the origins of antagonism in the context of a single species, wherein simultaneous targeting of two distinct cellular processes mitigates the defects due to targeting of only one of the processes (Brochado *et al*., 2018). Our findings reveal a wide range of phenomena explainable by resource competition alone, warranting caution when interpreting phenomena that might otherwise be attributed to chemical transformations of antibiotics or other biological mechanisms. Moreover, our results motivate quantification of the competitive landscape of communities (Ho *et al*., 2022b), whereby the determination of generalists and specialists will help to predict treatment outcomes.

There are countless combinations of complex antibiotic cocktails (simultaneous drug administration) and treatment scheduling strategies (such as an alternating antibiotic schedule (Marrec and Bitbol, 2020)) involving broad-spectrum antibiotics. To predict microbial community dynamics in response to such complex antibiotic perturbations, we analyzed two key “building block” scenarios at the heart of antibiotic treatment regimens: the swapping of application order of two narrow-spectrum antibiotics and the simultaneous application of two narrow-spectrum antibiotics. Understanding the interplay between resource competition and antibiotics in these minimal scenarios is the first step in predicting community dynamics during more complex antibiotic perturbations (and in communities with more species and nutrients) that occur in a clinical setting. For example, we can straightforwardly apply our model predictions during the simultaneous application of two narrow-spectrum antibiotics that each target a different species to a broad-spectrum antibiotic that simultaneously targets the two species.

Beyond responses of the human gut microbiota to antibiotics, our analyses can be broadly applied to other communities and to understand the effects of other environmental perturbations. Inhibition of growth is prevalent in other contexts, including soil communities, which produce numerous antimicrobials (Olanrewaju and Babalola, 2019), and wastewater communities, which can contain a wide spectrum of antibiotics (Jendrzejewska and Karwowska, 2018). In the context of our model, by allowing death rates to be both species-specific and also dependent on the presence of other species, our framework could be extended to scenarios in which antibiotics are produced and/or modified by community members. Moreover, microbial communities including the gut microbiota contain diverse bacteriophages whose deleterious effects on their bacterial host communities are not fully understood (Salmond and Fineran, 2015; Shkoporov et al., 2018). Changes in environmental parameters like temperature (Knapp and Huang, 2022), osmolarity (Tropini et al., 2018), or pH (Amor et al., 2020), likely rescale consumption rates in species-dependent manners similar qualitatively to antibiotics. *In vitro* measurements of the sensitivities of each community member to such parameters could be combined with parameterization of the competitive landscape (Ho *et al*., 2022b) to enable quantitative prediction of community responses.

In addition to killing, our findings suggest that deleterious agents can be used to engineer community dynamics at sublethal doses. Developing such engineering strategies will require new experiments to characterize the interplay between resource competition and antibiotic activity; these experiments include screens of communities grown *in vitro* (Aranda-Diaz et al., 2022; Goldford et al., 2018), and mouse models subject to lower than usual doses of antibiotics. It will also be important to analyze model behavior out of steady state, including serial dilution protocols mimicking the periodic turnover of the gut environment that could result in fundamentally different and interesting behaviors. Other interspecies interactions such as cross-feeding, which can be incorporated into CR models (Lopez and Wingreen, 2022), may also generate the potential for community-dependent antibiotic responses. This work provides a null model for these critical future endeavors.

## METHODS

### Simulations of population dynamics

We simulated species abundances over time in Python using the SciPy function scipy.integrate.solve_ivp, an explicit Runge-Kutta method for solving ordinary differential equations. We integrated all species abundances *n_i_* until they reached steady state, defined as when all species either satisfy 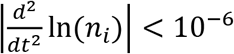 or *n_i_* decreases below a threshold abundance 10^-7^ (signifying extinction and removal from the community). To calculate richness, we counted the number of species that satisfied the conditions *n_i_* > 10^-7^ and 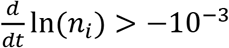 at steady state. For three species and three resources, the steady-state abundances of the three species can be directly calculated using a straightforward algorithm (Supplemental Text).

### Dimensionless parameters

For simulating population dynamics, we introduced dimensionless parameters for time *t′* ≡ *td*, for species abundance 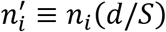, for resource consumption rate 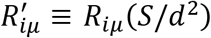, for death rate (*d* + *d_i_*)′ ≡ (*d* + *d_i_*)/*d*, and for resource supply rate 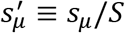. We can write Eq. 3 in terms of these dimensionless parameters as follows:

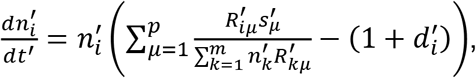

where 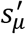 is the relative supply rate of resource *μ* such that 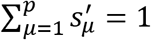.

### Simulations of randomly drawn communities

Unless stated otherwise, we selected 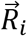, 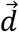 from a uniform distribution on the unit simplex, i.e., from the space of *m*-dimensional vectors such that the sum of the elements is 1. We chose 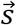 such that all resources were supplied at equal rates.

## ACKNOWLEDGEMENTS

We thank members of the Huang lab for helpful discussions. This work was funded by a Stanford Bioengineering Summer REU fellowship (to D.P.N.), a Stanford School of Medicine Dean’s Postdoctoral Fellowship (to P.H.), NIH Postdoctoral Fellowship F32 GM143859 (to P.H.), NSF Award EF-2125383 (to K.C.H.), and NIH Awards R01 AI147023 and RM1 GM135102 (to K.C.H.). K.C.H. is a Chan Zuckerberg Biohub Investigator.

## SUPPLEMENTAL FIGURES

**Figure S1:**
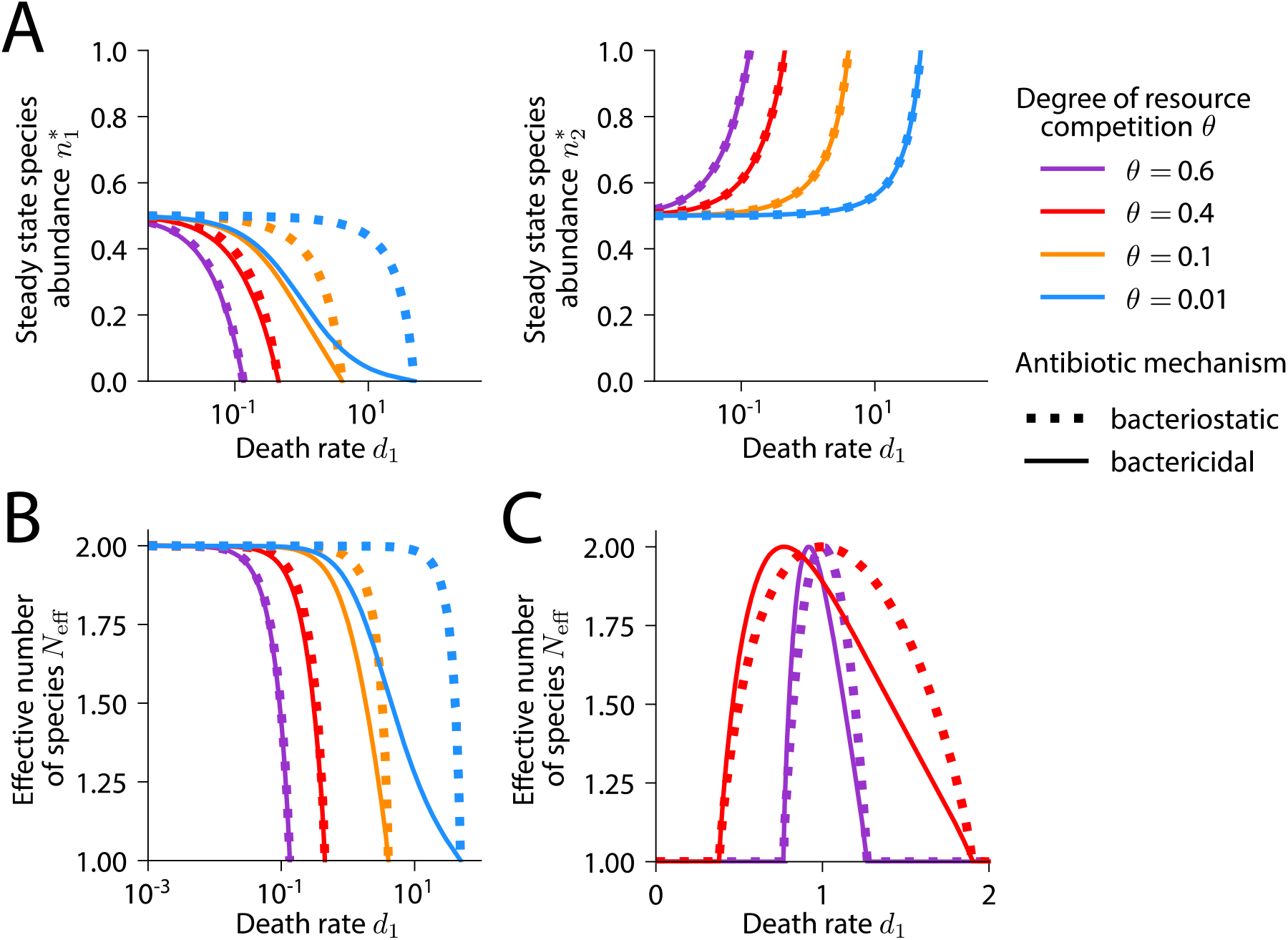
Bactericidal and bacteriostatic antibiotics have similar effects on species abundances. A) Simulations of a community with *m* = *p* = 2 defined by the resource consumption rate matrix *R* = ((1,0), (0,1)) > where *θ* parametrizes the niche overlap between the two species. Species 1 was targeted with increasing death rate *d*_1_. (Left) The steady state abundance of species 1 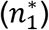 during treatment with a bactericidal antibiotic was always lower in comparison with the corresponding perturbation with a bacteriostatic antibiotic, given by *b*_1_ = (*d* + *d*_1_)/*d* (Supplemental Text). (Right) The steady-state abundance of species 2 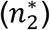 was independent of the antibiotic mechanism. B) For the community in (A), *N*_eff_ was generally higher at steady state for bacteriostatic versus bactericidal antibiotics. C) For a community defined by the resource consumption rate matrix *R* = ((2, 2*θ*),(*Θ*, 1)), steady-state evenness *N*_eff_ was non-monotonic (Supplemental Text), analogous to the richness in Fig. 2A. The difference in *N*_eff_ between a bacteriostatic and bactericidal antibiotic was greater when there was less niche overlap (*θ* = 0.4).

**Figure S2:**
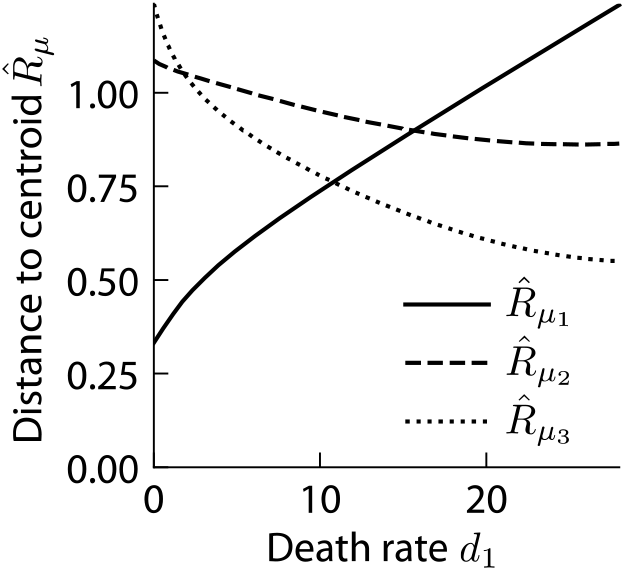
The centroid of the coexistence region shifts towards the niche of the targeted species during antibiotic treatment. For the community in Fig. 3A, we plotted the distance between the centroid of the coexistence region and each corner of the simplex, corresponding to each consumption niche. The niche of the targeted species (blue) is resources 2 and 3, so targeting the blue species caused the non-targeted species to encroach on the niche of the targeted species, as indicated by the centroid of the coexistence region moving closer to the corners of the simplex corresponding to resources 2 and 3, and farther from resource 1.

**Figure S3:**
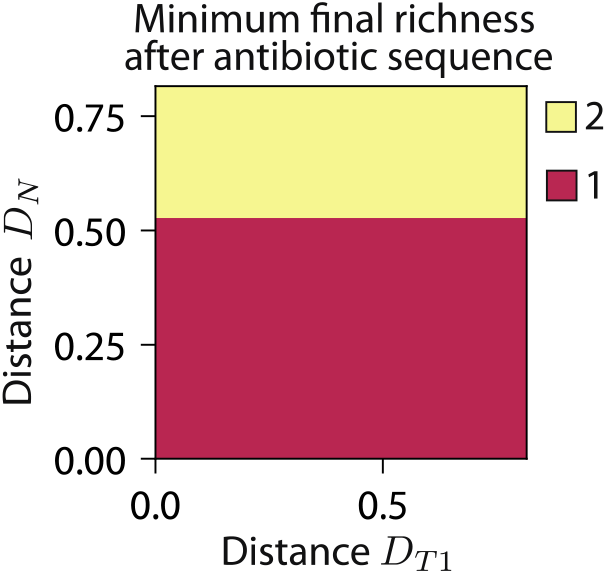
Final richness after antibiotic sequencing is always ≥ 2 when ≳ 0.5, precluding non-transitivity. We calculated the final richness after all antibiotic sequences (and reverse sequences) across the parameters (*D*_*T*1_, *D*_*T*2_, *D_N_, d*_1_, *d*_2_). The minimum final richness after all antibiotic sequences with *D_N_* ≳ 0.5 was 2. For Δ*ρ* to be nonzero (non-transitivity), the final richness after an antibiotic sequence must be 1, indicating that non-transitivity is not possible when *D_N_* ≳ 0.5.

**Figure S4:**
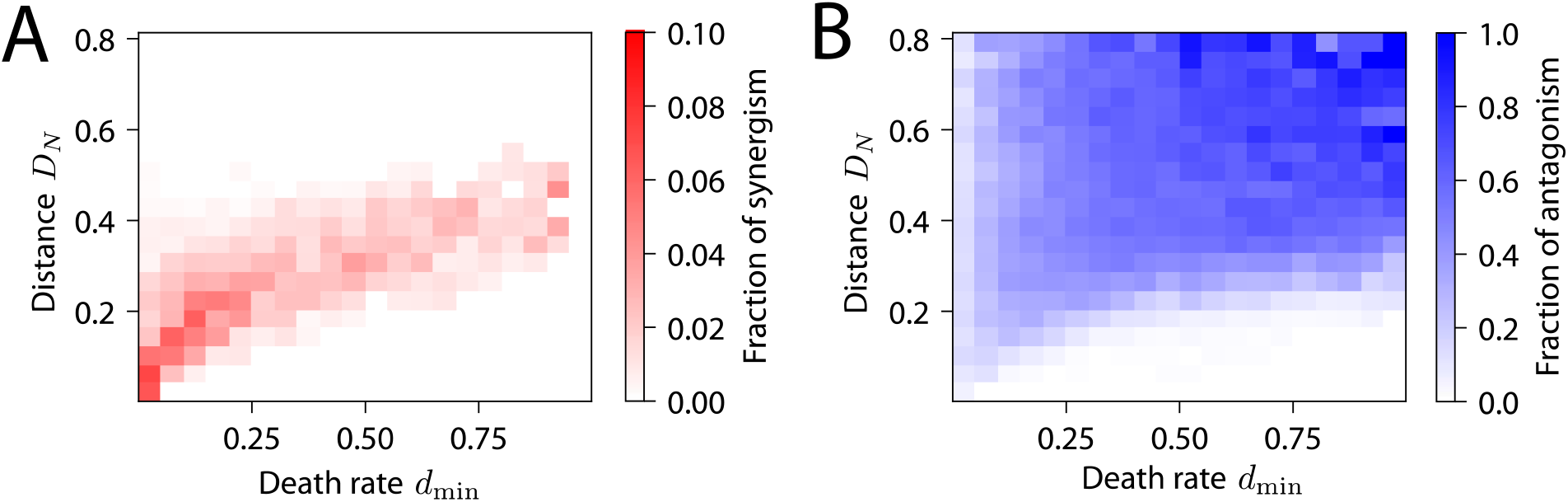
Non-additivity requires a balance between *D_N_* and death rates. A) For communities with higher *D_N_*, representing a more specialized non-targeted species, the fraction exhibiting synergism was maximal at larger values of *d*_min_ (the smaller of the two death rates). B) Communities with higher *d*_min_, the fraction exhibiting antagonism increased with increasing *D_N_*. These simulations generalize the results in Fig. 5H in the main text to communities without symmetry in consumption rates.

