## Supplemental Information for "Antibiotic effects on microbial communities are modulated by resource competition"

### Supplementary Information for "Antibiotic effects on microbial communities are modulated by resource competition"

#### Model details

We build on the consumer-resource (CR) model in (Posfai et al., 2017), which presents a model for $m$ species competing for $p$ types of steadily supplied resources at rates $\vec{s} = (s_1, \ldots, s_p)$ where $\sum_{\mu=1}^{p} s_\mu = S$ and $s_\mu$ is the supply rate of resource $\mu$. The dilution rate of the chemostat is $d$ but the resource loss rates (due to efflux from the chemostat) are approximated to be zero; this approximation simplifies calculations and has been shown to not significantly affect population dynamics (Posfai et al, 2017). Species $i$ has resource-consumption rates $\vec{R}_i = (R_{i1}, \ldots, R_{ip})$, and enzyme budget $E_i = \sum_{\mu=1}^{p} R_{i\mu}$, where $R_{i\mu}$ is the rate at which species $i$ consumes resource $\mu$. As stated in the main text, we begin with a classic formulation of a well-mixed CR model in a chemostat with dilution rate $d$ (Posfai et al., 2017),

$$\frac{dn_i}{dt} = (g_i(\vec{n}) - d)n_i, \quad (1)$$

where $n_i$ is the abundance of species $i$, $\vec{n} = (n_1, \ldots, n_m)$ is the vector of species abundances, and $g_i$ is the growth rate of species $i$, given by

$$g_i(\vec{n}) = \sum_{\mu=1}^{p} R_{i\mu} c_\mu, \quad (2)$$

where $c_\mu$ is the concentration of resource $\mu$:

$$c_\mu = \frac{s_\mu}{\sum_{k=1}^{m} n_k R_{k\mu}}. \qquad (3)$$

Combining Eq. 1-3 results in a single equation governing the population dynamics when all species have uniform 'death' rate $d$ (due to dilution):

$$\frac{dn_i}{dt} = n_i \left( \sum_{\mu=1}^{p} \frac{R_{i\mu} s_\mu}{\sum_{k=1}^{m} n_k R_{k\mu}} - d \right). \qquad (4)$$

#### Introduction of species-specific death rates $d_i$ to model perturbation by bactericidal antibiotics

To model bacteriostatic antibiotics, we assumed that the consumption rates $R_{i\mu}$ of species $i$ decrease by a factor $b_i$, leading to the following modification of Eq. 4:

$$\frac{dn_i}{dt} = n_i \left( \sum_{\mu=1}^{p} \frac{(R_{i\mu}/b_i) s_\mu}{\sum_{k=1}^{m} n_k (R_{k\mu}/b_k)} - d \right). \quad (5)$$

To model the effects of bactericidal antibiotics, we assumed that species $i$ experiences death at rate $d_i$ in addition to the effects of dilution, leading to the following modification of Eq. 4 to include both the dilution rate and the species-specific death rate $d_i$:

$$\frac{dn_i}{dt} = n_i\left(\sum_{\mu=1}^{p} \frac{R_{i\mu}s_\mu}{\sum_{k=1}^{m} n_k R_{k\mu}} - (d + d_i)\right). \qquad (6)$$

Note that throughout the text, we refer to the death rate as the sum of the dilution rate and the species-specific death rate. The two seemingly different modifications to the CR model in Eq. 5 and 6 can be related by a transformation of the consumption rates $R'_{i\mu} = R_{i\mu}/b_i$, where $b_i = 1 + d_i/d$, as well as a change of variables $n_i \equiv n'_i/b_i$. Plugging this change of variables into Eq. 6,

$$\frac{d(n'_i/b_i)}{dt} = \frac{n'_i}{b_i}\left(\sum_{\mu=1}^{p} \frac{R_{i\mu}s_\mu}{\sum_{k=1}^{m} n'_k R_{k\mu}/b_k} - (d + d_i)\right),$$

from which it follows that,

$$\frac{dn'_i}{d\tau_i} = n'_i\left(\sum_{\mu=1}^{p} \frac{R'_{i\mu}s_\mu}{\sum_{k=1}^{m} n'_k R'_{k\mu}} - d\right) \qquad (7)$$

for time rescaling $\tau_i \equiv tb_i$ and scaling of the resource consumption rates $R'_{i\mu} \equiv R_{i\mu}/b_i$. The solutions to Eq. 5 and Eq. 7 are identical, up to a scaling of the amplitudes ($n'_i \equiv n_i b_i$) and a scaling of time ($\tau \equiv tb_i$). Therefore, since $b_i \neq 0$, a set of species will coexist in the scenario governed by Eq. 5 if and only if that set of species coexists in the scenario governed by Eq. 7.

To summarize, consider a set $\mathcal{C}$ of $p$ supplied resources and $m$ species with resource consumption rates $R_{i\mu}$ and species-specific death rates $d_i + d$. Consider the transformation $\mathcal{C} \mapsto \mathcal{C}'$ defined by a transformation of the consumption rates and death rates,

$$R_{i\mu} \mapsto R'_{i\mu} = \frac{dR_{i\mu}}{d+d_i} \text{ and } d_i + d \mapsto d. \, (8)$$

This transformation preserves the property of coexistence, i.e., species $i \in \mathcal{C}$ coexists ($n_i^* > 0$) if and only if species $i \in \mathcal{C}'$ coexists. In other words, for a community with uniform death rates $d$ in steady state that is perturbed by bactericidal antibiotics (i.e., the death rate of species $i$ becomes $d + d_i$), we can view this perturbation as simply a transformation of the species $i$ enzyme budget of $E_i \mapsto E_i' = dE_i/(d+d_i)$ when considering the effect of the perturbation on coexistence. We therefore consider communities with uniform death rate $d$ when proving the coexistence rules because these rules can apply to communities with species-specific death rates using the transformation given by Eq. 8.

#### Proof of coexistence rules for $\boldsymbol{m}$ species, $\boldsymbol{p}$ resources, arbitrary enzyme budgets, and uniform death rates $\boldsymbol{d}$

**Lemma 1:** *If the consumption rates of two species $A$ and $B$ are linearly dependent, i.e., for all resources $\mu$, $R_{A\mu} = cR_{B\mu}$ for some positive constant $c \neq 1$, then species $A$ and $B$ cannot coexist if they have equal death rates $d$.*

*Proof:* For species $A$ and $B$ to coexist, their steady state growth rates $g_A$ and $g_B$, respectively, must be equal to their death rates $d_A = d$ and $d_B = d$, respectively. However, this equality cannot occur since $g_A$ cannot equal $g_B$ due to the relation $g_A = \sum_{\mu=1}^{p} R_{A\mu} c_\mu = \sum_{\mu=1}^{p} cR_{B\mu} c_\mu = cg_B \neq g_B$. Therefore, in what follows, we assume no pair of species strategies are linearly dependent.

**Lemma 2:** *Species $A$ and $B$ with equal death rate $d$ cannot coexist if $R_{A\mu} > R_{B\mu}$ for all resources $\mu$.*

*Proof:* Consider the growth rate of species $A$ and $B$ to be $g_A$ and $g_B$, respectively. For the two species to coexist, $g_A$ and $g_B$ must equal the death rate $d$ at steady state. However, $g_A$ cannot equal $g_B$, because $g_A = \sum_{\mu=1}^{p} R_{A\mu} c_\mu > \sum_{\mu=1}^{p} R_{B\mu} c_\mu = g_B$ since the resource concentrations $c_\mu$ must be positive. Therefore, in what follows, we assume no pair of species $A$ and $B$ satisfies $R_{A\mu} > R_{B\mu}$ for all resources μ.

**Lemma 3:** *The resource consumption rates* $R_{i\mu}$ *of a set of coexisting species with equal death rates* $d$ *must obey a linear metabolic tradeoff whereby there exist steady state nutrient concentrations* $c_\mu^* > 0$ *such that* $\sum_{\mu=1}^{p} R_{i\mu}\, c_\mu^* = d$ *for all species* $i$.

*Proof:* Since the supply rate of all resources is positive by assumption, by Eq. 3, the nutrient concentrations $c_\mu$ must always be positive, from which it follows that the steady-state nutrient concentrations $c_\mu^*$ must be positive. Furthermore, by Eq. 1, all species coexist when the steady state growth rates $g_i^*$ satisfy $g_i^* = d$. Thus, by Eq. 2, there must exist steady-state resource concentrations $c_\mu^* > 0$ such that $g_i^* = \sum_{\mu=1}^{p} R_{i\mu} c_\mu^* = d$ if the set of species coexist.

We can interpret this coexistence condition as a geometric constraint on the resource consumption strategy vectors $\vec{R}_i$ in the space of resource consumption rates as follows. If all species coexist, then since $\sum_{\mu=1}^{p} R_{i\mu} c_\mu^* = d$, all resource consumption strategies $\vec{R}_i$ lie on the $p$-dimensional hyperplane in the space of resource consumption rates defined by its axis intercepts, where the intercept of the axis corresponding to resource $\mu$ is $d/c_\mu^*$.

**Lemma 4:** *For* $p$ *resources and* $m$ *species with equal death rates* $d$*, all species coexist if and only if there exist* $c_\mu^* > 0$ *such that* $\sum_{\mu=1}^{p} R_{i\mu}\, c_\mu^* = d$ *for all species* $i$ *and the normalized supply rate vector* $\hat{s} \equiv (d/S)\vec{s}$ *lies in the convex hull of the rescaled consumption strategies* $\vec{\hat{R}}_i = \left(c_1^* R_{i1}, \dots, c_p^* R_{ip}\right)$.

*Proof:* There are two cases: $m \geq p$ and $m < p$, and we consider each separately. The statement was proved for the case $m \geq p$ in (Posfai *et al.*, 2017). Here, we prove the statement for the other case, $m < p$; our proof holds for $m = p$ as well.

For coexistence of all species in the community, we need $dn_i/dt = 0$ at steady-state abundances $n_i^* > 0$, so by Eq. 1, this requirement is equivalent to $g_i(\vec{n}^*) = d$, or

$$d = \sum_{\mu=1}^{p} R_{i\mu}\, c_\mu^*, \qquad (9)$$

where

$$c_\mu^* = \frac{s_\mu}{\sum_{k=1}^m n_k^* R_{k\mu}}. \qquad (10)$$

For coexistence of all species, we need a solution $n_i^* > 0$ and $c_\mu^* > 0$. Since Eq. 9 must hold for all species $i$, it represents $m$ equations, and since all resource consumption strategies $\vec{R}_i$ are linearly independent, we are left with $p - m$ free variables. Rewriting $c_\mu^*$ in terms of the $p - m$ free variables and plugging into Eq. 10, we now have $p$ unknowns since we must solve for the $m$ values of $n_i^*$ and the $p - m$ free variables from Eq. 9. Since Eq. 10 represents $p$ distinct equations due to the linear independence of $\vec{R}_i$, there are two cases: there is a unique solution for the values of $n_i^* > 0$ and $c_\mu^* > 0$, or there is not. Note that we used the linear independence of $\vec{R}_i$ and the fact that we have a total of $m + p$ distinct equations and $m + p$ unknowns to rule out the possibility that there are an infinite number of solutions.

Defining the rescaled consumption rate vector $\vec{\hat{R}}_i = \left(c_1^* R_{i1}, \dots, c_p^* R_{ip}\right)$ and rearranging Eq. 10, in vector notation, we conclude that all species coexist if the system of equations

$$n_1^* \vec{\hat{R}}_1 + \cdots + n_m^* \vec{\hat{R}}_m = \vec{s}, \qquad (11)$$

has a positive solution $n_i^* > 0$. It follows from Eq. 9 and 10 that $\sum_{k=1}^m n_k^* = S/d$, so we define the normalized supply rate vector $\hat{s} \equiv (d/S)\vec{s}$ and observe that Eq. 11 having a positive solution is equivalent to the following equation having a positive solution:

$$n_1^* \vec{\hat{R}}_1 + \cdots + n_m^* \vec{\hat{R}}_m = \hat{s}, \text{ where } \sum_{i=1}^m n_i^* = 1, \qquad (12)$$

which is equivalent to $\hat{s}$ lying in the convex hull of the vectors $\vec{\hat{R}}_i$. Note that the convex hull normally includes nonnegative solutions $n_i^* \geq 0$ of Eq. 12, but we assume that no values of $n_i^*$ are zero, i.e., we assume that $\hat{s}$ does not lie on the perimeter of the convex hull of the vectors $\vec{\hat{R}}_i$.

We have shown that there exists a unique steady-state solution $n_i^* > 0$ if and only if $\hat{s}$ lies in the convex hull of the vectors $\vec{\hat{R}}_i$ for positive steady-state nutrient consumption rates satisfying $d = \sum_{\mu=1}^{p} R_{i\mu} c_\mu^*$. We next show that the unique steady state solution $n_i^*$ is an attractor of the system.

As in (Posfai *et al.*, 2017), we write $\vec{n} = \vec{n}^* + \Delta\vec{n}$ and linearize the growth equations in Eq. 1-3 around the unique fixed point $\vec{n}^*$:

$$\frac{d\Delta n_i}{dt} \approx \left(\sum_{\tau=1}^{m} \frac{\partial g_i}{\partial n_\tau}(\vec{n}^*)\Delta n_\tau\right) n_i^*. \qquad (13)$$

Assuming coexistence, and therefore a unique steady state solution $n_i^*$, $c_\mu^*$, we can treat these as known constants that do not depend on $\Delta\vec{n}$. We next calculate the partial derivatives of the growth rate at $\vec{n}^*$,

$$\frac{\partial g_i}{\partial n_\tau}(\vec{n}^*) = -\sum_{\mu=1}^{p} R_{i\mu} R_{\tau\mu} \frac{s_\mu}{\left(\sum_{k=1}^{m} n_k^* R_{k\mu}\right)^2},$$

then plug into Eq. 13 to obtain the linearized system of growth equations,

$$\left(\frac{d\Delta n_1}{dt}, \dots, \frac{d\Delta n_m}{dt}\right)^T = -DM(\Delta n_1, \dots, \Delta n_m)^T,$$

where $D \equiv \mathrm{diag}(\vec{n}^*)$, and $M_{ij} \equiv \sum_{\mu=1}^{p} R_{i\mu} R_{j\mu} \frac{s_\mu}{\left(\sum_{k=1}^{m} n_k^* R_{k\mu}\right)^2}$. We next argue that the eigenvalues of $DM$ are nonnegative, from which it follows that the unique solution $n_i^* > 0, c_\mu^*$ is an attractor of the system.

Here, we use an argument similar to (Posfai *et al.*, 2017). Since $n_i^* > 0$, $D$ is invertible, so the transformation $DM \mapsto D^{1/2} M D^{1/2}$ is a similarity transformation, from which it follows that the eigenvalues of $D^{1/2} M D^{1/2}$ are identical to those of $DM$. For arbitrary vector $\vec{v}$,

$$\vec{v}^T D^{1/2} M D^{1/2} \vec{v} = \sum_{\mu=1}^{p} \frac{\left(\sum_{k=1}^{m} n_k^* R_{k\mu}\right)^2}{s_\mu} \left(\sum_{i=1}^{m} v_i R_{i\mu} \sqrt{n_i^*}\right) \left(\sum_{j=1}^{m} v_j R_{j\mu} \sqrt{n_j^*}\right) =$$

$$\sum_{\mu=1}^{p} \frac{\left(\sum_{k=1}^{m} n_k^* R_{k\mu}\right)^2}{s_\mu} \left(\sum_{i=1}^{m} v_i R_{i\mu} \sqrt{n_i^*}\right)^2 \geq 0,$$

so $D^{1/2} M D^{1/2}$ is symmetric and positive semi-definite, from which it follows that $D^{1/2} M D^{1/2}$ and thus $DM$ has nonnegative eigenvalues.

Using Lemma 4, we can determine whether all species coexist in a community of $p$ resources and $m$ species with arbitrary enzyme budgets $E_i = \sum_{\mu=1}^{p} R_{i\mu}$ and equal death rates $d$. However, in the case that not all species in the community $\mathcal{C}$ coexist, we can determine which subset $\mathcal{S} \subseteq \mathcal{C}$ of species coexists as follows.

A subset of species $\mathcal{S} \subseteq \mathcal{C}$ will coexist if there exist steady-state nutrient concentrations $c_\mu^* > 0$ such that $\sum_{\mu=1}^{p} R_{i\mu}\, c_\mu^* = d$ and $\hat{s}$ lies in the convex hull of the vectors $\vec{R}_i$, and no other species $j \in \mathcal{C} - \mathcal{S}$ can invade the coexisting subset $\mathcal{S}$.

We consider whether species $j \in \mathcal{C} - \mathcal{S}$ can invade the subset $\mathcal{S}$ by adding species $j$ to the community of species in $\mathcal{S}$ at an infintesimal abundance $n_j \ll n_s$ for all $s \in \mathcal{S}$, and calculate $dn_j/dt$, the rate of change of abundance of species $j$. If the ratio $\frac{dn_j/dt}{n_j}$ is negative and is independent of $n_j$, then species $j$ goes extinct, and therefore cannot invade the community and coexist. If the ratio $\frac{dn_j/dt}{n_j}$ is positive, then species $j$ can invade the community and coexist.

**Lemma 5:** *Consider a set $\mathcal{C}$ of species with equal death rates $d$ that coexist with steady state resource concentrations $c_\mu^* > 0$ such that $\sum_{\mu=1}^{p} R_{i\mu} c_\mu^* = d$ for all species $i \in \mathcal{C}$. If a new community $\mathcal{C}' = \mathcal{C} \cup \{j\}$ does not satisfy the requirements for all species in $\mathcal{C}'$ to coexist, then species $j$ can invade the set of species in $\mathcal{C}$ if $\sum_{\mu=1}^{p} R_{i\mu}\, c_\mu^* > d$, and species $j$ cannot invade the set of species in $\mathcal{C}$ if $\sum_{\mu=1}^{p} R_{i\mu}\, c_\mu^* < d$.*

*Proof:* Consider a set $\mathcal{C}$ of $m$ coexisting species with $p$ resources, i.e., there exist $c_\mu^* > 0$ such that $\sum_{\mu=1}^{p} R_{i\mu} c_\mu^* = d$ for all species $i \in \mathcal{C}$. Next, consider adding another species $j$ creating a new set $\mathcal{C}' = \mathcal{C} \cup \{j\}$, where the set of species $\mathcal{C}'$ does not satisfy the requirements for all species in $\mathcal{C}'$ to coexist. Therefore, we know that at least one species in $\mathcal{C}'$ must go extinct. To determine if species $j \in \mathcal{C}'$ goes extinct, we calculate $dn_j/dt$ when $n_j$ is infinitesimal such that for all other species $i \in \mathcal{C}$, $n_i$ are the steady state abundances just before species $j$ was introduced. We first calculate the perturbed resource concentration $c_\mu$ to first order in $\epsilon_\mu \equiv n_j R_{j\mu} c_\mu^*/s_\mu \ll 1$,

$$c_\mu = \left(\frac{n_j R_{j\mu}}{s_\mu} + \frac{\Sigma_{k\in\mathcal{C}} n_k R_{k\mu}}{s_\mu}\right)^{-1} = \left(\frac{\epsilon_\mu}{c_\mu^*} + \frac{1}{c_\mu^*}\right)^{-1} \approx c_\mu^*\left(1-\epsilon_\mu\right), \quad (14)$$

where $c_\mu^*$ is the concentration of resource $\mu$ when community $\mathcal{C}$ coexists at steady state, so that we can use Eq. 10. Plugging in Eq. 14 to calculate the approximate growth rate of species $i$,

$$g_i(\vec{n}) \approx \sum_{\mu=1}^{p} R_{i\mu}\, c_\mu^*\left(1-\epsilon_\mu\right),$$

which is valid for $i \in \mathcal{C}$ or $i = j$. To determine whether species $j$ can invade the community $\mathcal{C}$, we first use Eq. 1 and the above approximation to calculate the (approximate) ratio $\frac{dn_i/dt}{n_i}$ for $i \in \mathcal{C}$:

$$\frac{dn_i/dt}{n_i} \approx \sum_{\mu=1}^{p} R_{i\mu}\, c_\mu^*\left(1-\epsilon_\mu\right) - d = -\sum_{\mu=1}^{p}\left(R_{i\mu} c_\mu^*\right)\epsilon_\mu,$$

since $\sum_{\mu=1}^{p} R_{i\mu} c_\mu^* = d$. Next, we similarly calculate the same (approximate) ratio for species $j$,

$$\frac{dn_j/dt}{n_j} \approx \sum_{\mu=1}^{p} R_{i\mu}\, c_\mu^*\left(1-\epsilon_\mu\right) - d \approx \sum_{\mu=1}^{p} R_{i\mu} c_\mu^* - d, \qquad (15)$$

since $\sum_{\mu=1}^{p} R_{i\mu} c_\mu^* \neq d$. We analyze Eq. 15 separately for the two cases $\sum_{\mu=1}^{p} R_{j\mu}\, c_\mu^* < d$ and $\sum_{\mu=1}^{p} R_{j\mu}\, c_\mu^* > d$.

**Case $\sum_{\mu=1}^{p} R_{j\mu}\, c_\mu^* < d$:** Here, the ratio $\frac{dn_j/dt}{n_j}$ is negative and independent of $n_j$ (i.e., order zero in $\epsilon_\mu$), and the ratio $\frac{dn_i/dt}{n_i}$ for $i \in \mathcal{C}$ is order $\epsilon_\mu \ll 1$. Therefore, species $j$ cannot invade community $\mathcal{C}$, i.e. species $j$ goes extinct, leaving all species in $\mathcal{C}$ able to coexist.

**Case $\sum_{\mu=1}^{p} R_{j\mu}\, c_\mu^* > d$:** Here, the ratio $\frac{dn_j/dt}{n_j}$ is positive and order zero in $\epsilon_\mu$, and the ratio $\frac{dn_i/dt}{n_i}$ for $i \in \mathcal{C}$ is negative and order $\epsilon_\mu \ll 1$. Therefore, species $j$ can invade community $\mathcal{C}$, meaning that at least one species in $\mathcal{C}$ must go extinct, since $\mathcal{C} \cup \{j\}$ does not satisfy the condition for coexistence by assumption.

By combining Lemmas 3-5, we obtain the coexistence rules presented in the main text.

#### Closed-form solution for steady-state abundances of $m$ coexisting species when competing for $p = m$ resources

When the $m$ species satisfy the convex hull coexistence condition, we can solve for the unique steady state abundances $\vec{n}^*$. Starting from Eq. 6, we set the growth rate $g_i$ equal to the death term $d + d_i$ for each species and write the equations in matrix form,

$$\overrightarrow{n^*} RC = \vec{s} \qquad (16)$$

$$R\vec{c}^T = \vec{d}^T \qquad (17)$$

where $R = \{R_{i\mu}\}$, $\vec{d} = (d + d_1, \ldots, d + d_m)$, $C = \text{diag}(c_\mu^*)$, and $\vec{c} = (c_1^*, \ldots, c_p^*)$. $R$ is a square matrix since $m = p$, and all species have linearly independent consumption strategies so the rows of $R$ are linearly independent, therefore $R$ is invertible. Since the steady-state nutrient concentrations are positive and all resources are supplied at positive rates, the diagonal matrix $C$ is positive along the diagonal, thus $C$ is invertible. Therefore, we rearrange Eq. 16 and 17 as follows:

$$\vec{n}^* = \vec{s} C^{-1} R^{-1} \qquad (18)$$

where the diagonal elements of $C$ are given by

$$\vec{c} = R^{-1}\vec{d}^T. \qquad (19)$$

#### Comparison of steady-state abundances for bactericidal versus bacteriostatic antibiotics when $m = p = 2$

By comparing the population dynamics with species-specific death rates $d_i$ in Eq. 9 and species-specific resource consumption rate reduction factors $b_i$ in Eq. 5, the steady-state abundances are not in general the same in these two scenarios despite the set of coexisting species being identical when $b_i = (d + d_i)/d$.

When $m = p = 2$, we can solve explicitly for the steady state abundances $\vec{n}^*$ using Eq. 18 and 19. We focus on a resource consumption rate matrix $R = \big((1, \theta), (\theta, 1)\big)$, dilution rate $d = 1$, and equal supply rates for both resources $\vec{s} = (0.5, 0.5)$.

*Bactericidal antibiotics*

We consider varying the death rate of species 1 $d_1$ from 0 to $d_1^{\max}$ while keeping the death rate of species 2 at zero. The death rate vector is therefore $\vec{d} = (d_1 + 1{,}1)$ since the dilution rate $d$ is set to 1. Plugging into Eq. 18 and 19, the steady state abundances are

$$\vec{n}^*_{\text{cidal}} = \frac{1}{2}\left(\frac{1}{d_1+1-\theta} + \frac{\theta}{\theta+d_1\theta-1}, \frac{1}{1-d_1\theta-\theta} + \frac{\theta}{\theta-d_1-1}\right). \quad (20)$$

We can solve for the maximum death rate of species 1 $d_1^{\max}$, the death rate above which species 1 goes extinct, by solving for the death rate such that the steady-state abundance of species 1 $n^*_{1,\text{cidal}}$ is zero:

$$d_1^{\max} = \frac{\theta^2-2\theta+1}{2\theta}.$$

*Bacteriostatic antibiotics*

We next consider reducing the consumption rates of species 1 by a factor $b_1$, giving the new consumption rate matrix, $R = \big((1/b_1, \theta/b_1), (\theta, 1)\big)$, with zero death aside from the dilution of the chemostat so $\vec{d} = (1{,}1)$. To directly compare the bacteriostatic and bactericidal cases, we take $b_1 = (d + d_1)/d$, motivated by the fact that the coexistence is identical in bacteriostatic and bactericidal cases with this relationship between $b_1$ and $d_1$. Plugging into Eq. 18 and 19, the steady state abundances are

$$\vec{n}^*_{static} = \frac{1}{2}\left(\frac{d_1+1}{d_1+1-\theta} + \frac{\theta d_1+\theta}{\theta+d_1\theta-1}, \frac{1}{1-d_1\theta-\theta} + \frac{\theta}{\theta-d_1-1}\right). \quad (21)$$

By comparing Eq. 20 and 21, the steady state abundance of the non-targeted species 2 is the same in the bactericidal and bacteriostatic cases (i.e. $n^*_{2,\text{cidal}} = n^*_{2,\text{static}}$, Fig. S1A). However, for species 1, in the range $d_1 \in [0, d_1^{\max}]$,

$$n^*_{1,\text{static}} = \frac{1}{2}\left(\frac{d_1+1}{d_1+1-\theta} + \frac{\theta d_1+\theta}{\theta+d_1\theta-1}\right) = n^*_{1,\text{cidal}} + \frac{1}{2}\left(\frac{d_1}{d_1+1-\theta} + \frac{\theta d_1}{\theta+d_1\theta-1}\right) \geq n^*_{1,\text{cidal}},$$

where equality occurs when $d_1 = 0$ or $d_1^{\max}$. In other words, the community is more even at steady state during treatment with bacteriostatic antibiotics (Fig. S1A). Notably, the difference in steady-state abundance of the targeted species during bactericidal versus bacteriostatic antibiotics decreased when $\theta$ increased; larger $\theta$ corresponds to a larger degree of resource competition. This finding is consistent with the behavior at the extreme limits: when there is no resource competition, the steady-state abundances are unaffected by reducing the resource consumption rates for one species, but varying the death rate of one species must change the steady-state abundances regardless of the degree of resource competition.

##### Condition for maximizing the effective number of species at steady state $N_{\text{eff}}$

Consider $m$ species competing for $p = m$ resources. To maximize $N_{\text{eff}}$, we solve for the strategies $R_{i\mu}$ such that $dn_i/dt = 0$ at steady-state abundances $n_i^* = 1/N$, where $N = \sum_{i=i}^{p} n_i^*$. Plugging these conditions into Eq. 6 and defining the fractional consumption rate $F_{i\mu} \equiv R_{i\mu}/\sum_{k=1}^{m} R_{k\mu}$,

$$d + d_i = \sum_{\mu=1}^{p} \frac{R_{i\mu} s_\mu}{\Sigma_k n_k R_{k\mu}} = \frac{1}{N} \sum_{\mu=1}^{p} \frac{R_{i\mu} s_\mu}{\Sigma_k R_{k\mu}} = \frac{1}{N} \sum_{\mu=1}^{p} F_{i\mu} s_\mu.$$

When all resources are supplied at equal rates $s_\mu = 1/S$, the condition for maximizing $N_{\text{eff}}$ becomes $\sum_{j=\mu}^{p} F_{i\mu} = \delta_i$ where $\delta_i = (d + d_i)NS$ is a constant. When all species have uniform death rate, the condition for maximizing $N_{\text{eff}}$ becomes $\sum_{\mu=1}^{p} F_{i\mu} = F_i = F$, which can be interpreted simply as the total fractional consumption rate $F_i$ being constant for all species.

##### Application of the coexistence rules to calculate steady-state abundances when $\boldsymbol{m = p = 3}$

When $m = p = 3$, steady-state abundances can be calculated without simulating population dynamics and instead by applying coexistence rules. From Eq. 6, the steady-state abundances $n_i^*$ satisfy $S = n_1^*(d + d_1) + \cdots + n_m^*(d + d_m)$, from which it follows that if species 1 is the only existing species at steady state, then $n_1^* = S/(d + d_1)$, and if species 1 and 2 are the only coexisting species at steady state, then $n_2^* = \frac{S - n_1^*(d + d_1)}{d + d_2}$. Thus, we can simplify the calculation of steady-state abundances and avoid calculating the full trajectory of population dynamics by using the following algorithm to calculate steady-state abundances.

**1. Check if all species coexist:** We solve for the steady-state abundances using Eq. 18 and 19, and if all steady state abundances are positive then all species can coexist, satisfying the assumptions of Eq. 18 and 19. If Eq. 18 and 19 yield steady-state abundances that are not all positive, then at least one species must go extinct and therefore Eq. 18 and 19 are not valid.

**2. Check if exactly one species exists:** Species $i$ is the sole existing species if the other two species cannot invade species $i$ at steady state. Thus, we calculate the steady-state nutrient concentrations such that species $i$ is the sole existing species using Eq. 10, giving $c_\mu^* = s_\mu(d + d_i)/(SR_{i\mu})$, using the fact that $n_i^* = S/(d + d_i)$ in this case. Next, we use Eq. 3 to calculate $g_j$, the growth rate of species $j \neq i$, and if $g_j < d_j$ then $j$ cannot invade $i$; otherwise, $j$ can invade $i$. If there is a species $i$ such that all other species $j \neq i$ cannot invade $i$, then species $i$ is the only existing species at steady state and its abundance is $n_i^* = S/(d + d_i)$.

**3. Check if exactly two species coexist:** After ruling out that three or one species coexist, we know that two species must coexist. The pair of coexisting species must be mutually invasible, meaning for example that species 1 can invade species 2 and vice versa. We say that species 1 can invade species 2 if, when species 2 is in monoculture at steady state and then perturbed by the addition of species 1 at infinitesimal abundance, the growth rate of species 1 is positive. We check this mutual invasibility condition for all three pairs of species. For a pair of species that is mutually invasible (here taken to be species 1 and 2), we assume these are the two species that coexist and calculate their steady-state abundances using the relation $n_2^* = \frac{S - n_1^*(d+d_1)}{d+d_2}$ and Eq. 6 to solve for $n_1^*$ such that $dn_1/dt$ is zero. The outcome is an order 3 polynomial in $n_1^*$ with two roots of $n_1^*$ equal to zero or $S/(d + d_1)$. We solve for the third root (which must be between zero and $S/(d + d_1)$). After doing so, we use Eq. 3 to calculate the resource concentrations at steady state (assuming that the pair of species are the two that coexist), and test whether the third species cannot invade. If the third species cannot invade, then we have determined the two coexisting species and calculated their steady-state abundances. If the third species can invade, then these two are not the pair of coexisting species.
